# Differential Conservation and Loss of CR1 Retrotransposons in Squamates Reveals Lineage-Specific Genome Dynamics across Reptiles

**DOI:** 10.1101/2024.02.09.579686

**Authors:** Simone M. Gable, Nicholas Bushroe, Jasmine Mendez, Adam Wilson, Brendan Pinto, Tony Gamble, Marc Tollis

## Abstract

Transposable elements (TEs) are repetitive DNA sequences which create mutations and generate genetic diversity across the tree of life. In amniotic vertebrates, TEs have been mainly studied in mammals and birds, whose genomes generally display low TE diversity. Squamates (Order Squamata; ∼11,000 extant species of lizards and snakes) show as much variation in TE abundance and activity as they do in species and phenotypes. Despite this high TE activity, squamate genomes are remarkably uniform in size. We hypothesize that novel, lineage-specific dynamics have evolved over the course of squamate evolution to constrain genome size across the order. Thus, squamates may represent a prime model for investigations into TE diversity and evolution. To understand the interplay between TEs and host genomes, we analyzed the evolutionary history of the CR1 retrotransposon, a TE family found in most tetrapod genomes. We compared 113 squamate genomes to the genomes of turtles, crocodilians, and birds, and used ancestral state reconstruction to identify shifts in the rate of CR1 copy number evolution across reptiles. We analyzed the repeat landscapes of CR1 in squamate genomes and determined that shifts in the rate of CR1 copy number evolution are associated with lineage-specific variation in CR1 activity. We then used phylogenetic reconstruction of CR1 subfamilies across amniotes to reveal both recent and ancient CR1 subclades across the squamate tree of life. The patterns of CR1 evolution in squamates contrast other amniotes, suggesting key differences in how TEs interact with different host genomes and at different points across evolutionary history.

## Introduction

Transposable elements (TEs) are repetitive DNA sequences that are important contributors to genome organization and diversity in eukaryotes. TEs create mutations by mobilizing throughout the genome via transposition, generating genetic diversity that evolutionary forces can act on, in some cases forming the basis of new phenotypes (Kidwell and Lisch 2001). TEs also constitute between 10% and 80% of some vertebrate genomes (Jang et al. 2019), including at least 60% of the human genome (de Koning et al. 2011), and high TE abundance is linked to increased genome size (Kapusta et al. 2017; Blommaert 2020). Comparative genomic studies of TE dynamics across species shed light on the origins and maintenance of a near ubiquitous form of biological variation.

Across the genomes of mammals, birds, and reptiles (i.e., amniotes), the CR1 (Chicken repeat 1) autonomous non-terminal repeat (non-LTR) retrotransposon is one of the most abundant and active TE families (Chalopin et al. 2015; Galbraith et al. 2021). First described from the chicken genome (*Gallus gallus;* Stumph et al. 1984; International Chicken Genome Sequencing Consortium 2004), CR1 elements are similar to other vertebrate non-LTR retrotransposon families, such as L1 and L2, and are equipped with the machinery to “copy and paste” themselves into new genomic locations by retrotransposition using their own encoded proteins, including a reverse transcriptase (Ichiyanagi and Okada 2008). While CR1 is found only scarcely across mammalian genomes, which are instead dominated by L1 retrotransposons and non-autonomous retrotransposons (i.e., SINEs) (Chalopin et al. 2015), CR1 is the dominant repetitive element in the genomes of birds and reptiles (i.e., sauropsids), comprising ∼4% of most avian genomes and up to 18% of some reptile genomes (Shedlock 2006; Shaffer et al. 2013; Green et al. 2014; Chalopin et al. 2015; Pasquesi et al. 2018; Gable et al. 2023).

While CR1 sequences have been found in the genomes of species from all amniote groups, the abundance and evolutionary origins of CR1 within each of the amniote clades varies widely (Shedlock et al. 2006; Suh et al. 2015). In a given amniote genome, CR1 elements can be further divided into subfamilies, each with unique phylogenetic histories indicative of a pattern of diversification that occurred during the evolution of their host genomes (Suh et al. 2015). The genomes of birds, which generally lack TE abundance and diversity overall (Kapusta and Suh 2017; Sotero-Caio et al. 2017), contain relatively few CR1 elements, derived from sets of avian-specific CR1 subfamilies (Suh et al. 2015). Compared to birds, other archosaurian-line sauropsids such as turtles and crocodilians have many more CR1 copies in their genomes (Chalopin et al. 2015; Sotero-Caio et al. 2017). CR1 elements in turtle and crocodilian genomes are also derived from a far more diverse set of CR1 subfamilies than in bird genomes (Suh et al. 2015), suggesting an ancient origin of CR1 retrotransposons that stems from the genome of the ancestral amniote, followed by the subsequent loss of most CR1 diversity in birds (Suh et al. 2015; Galbraith et al. 2021).

Outside turtles and archosaurs, the remaining living reptiles are the lepidosaurs, which include ∼11,000 extant species of squamates (Order Squamata; i.e., lizards and snakes) and a single species of tuatara (*Sphenodon punctatus*, which constitutes the outgroup to squamates; (Uetz, P., and J. Hošek 2023)). The genome of the tuatara contains CR1 elements which cluster phylogenetically with the more early-branching CR1 subfamilies found in turtles, archosaurs, and mammals, suggesting ancient features of amniote genome evolution (Gemmell et al. 2020). Phylogenetic reconstructions of CR1 sequences in the genome of the green anole lizard (*A. carolinensis*) revealed the coexistence of numerous CR1 subfamilies (Novick et al. 2009), and CR1 subfamilies from anole and Burmese python (*Python bivitattus*) genomes were found to be nested deep within the amniote CR1 phylogeny (Suh et al. 2015). This suggests that the most ancient CR1 subfamilies otherwise shared across most amniotes have been lost in squamate genomes, as they have in birds. However, a lack of genome assemblies from representatives of many squamate lineages have only been made available very recently (Gable et al. 2023; Pinto et al. 2023; Card et al. 2023), limiting our ability to accurately model ancestral states in reptile genome evolution (Shedlock 2006).

Here, we analyzed the evolutionary history of the CR1 retrotransposon across amniote evolution using a wide taxonomic sampling of whole genome assemblies from 345 species of reptiles and birds, including 113 squamates. Our study focused on questions about the abundance, activity, and diversity of CR1 retrotransposons in the genomes of squamates and other amniotes. We identified contrasting patterns of loss and retention of CR1 throughout amniote evolution, particularly in squamates, signaling important differences in TE evolutionary dynamics across the vertebrate classes. We also demonstrate that the diversification of squamates, which spans a ∼200 million year fossil history and ∼11,000 extant species (Brownstein et al. 2023; Uetz, P., and J. Hošek 2023), was accompanied by rapid shifts in genome evolution, multiple episodes of CR1 expansion as well as loss in CR1 copy number – a dynamic history of TE evolution within the host lineage that differs greatly from birds and other reptiles. We suggest that squamates are a promising model for TE evolution, with the potential to shed light on key biological questions about the origins of genome size and structural variation in vertebrates.

## Results

### Variation in CR1 Copy Number across Squamates Compared to other Reptiles

We identified and annotated repeats in genome assemblies for 113 squamate species representing seven major clades (Iguania, n=17; Anguimorpha, n=8; Serpentes, n=59; Lacertoidea, n=15; Scincoidea, n=4; Gekkota, n=9; Dibamia n=1; Fig. 1), 30 turtle species representing all extant families of cryptodires and pleurodires (Shaffer et al. 2017; Gable et al. 2022), and 204 archosaur species including 200 birds and four crocodilians using de novo repeat detection, followed by classification based on repeat databases (Materials and Methods; Supplementary Table 1). The full results of all repeat types detected in all analyzed genomes are summarized in Supplementary Table 1. The CR1 family was the dominant element on average across all reptile genomes, including squamates. Our analyses calculated 10.2 Gbp (10,236,613,375 bp) of CR1 sequences across 113 squamate genomes, with CR1 accounting for ∼5.2% of the average squamate genome, and non-LTR retrotransposons in general occupying ∼15.9% of the average squamate genome (after filtering for nested elements; Supplementary Table 1).

**Figure 1:**
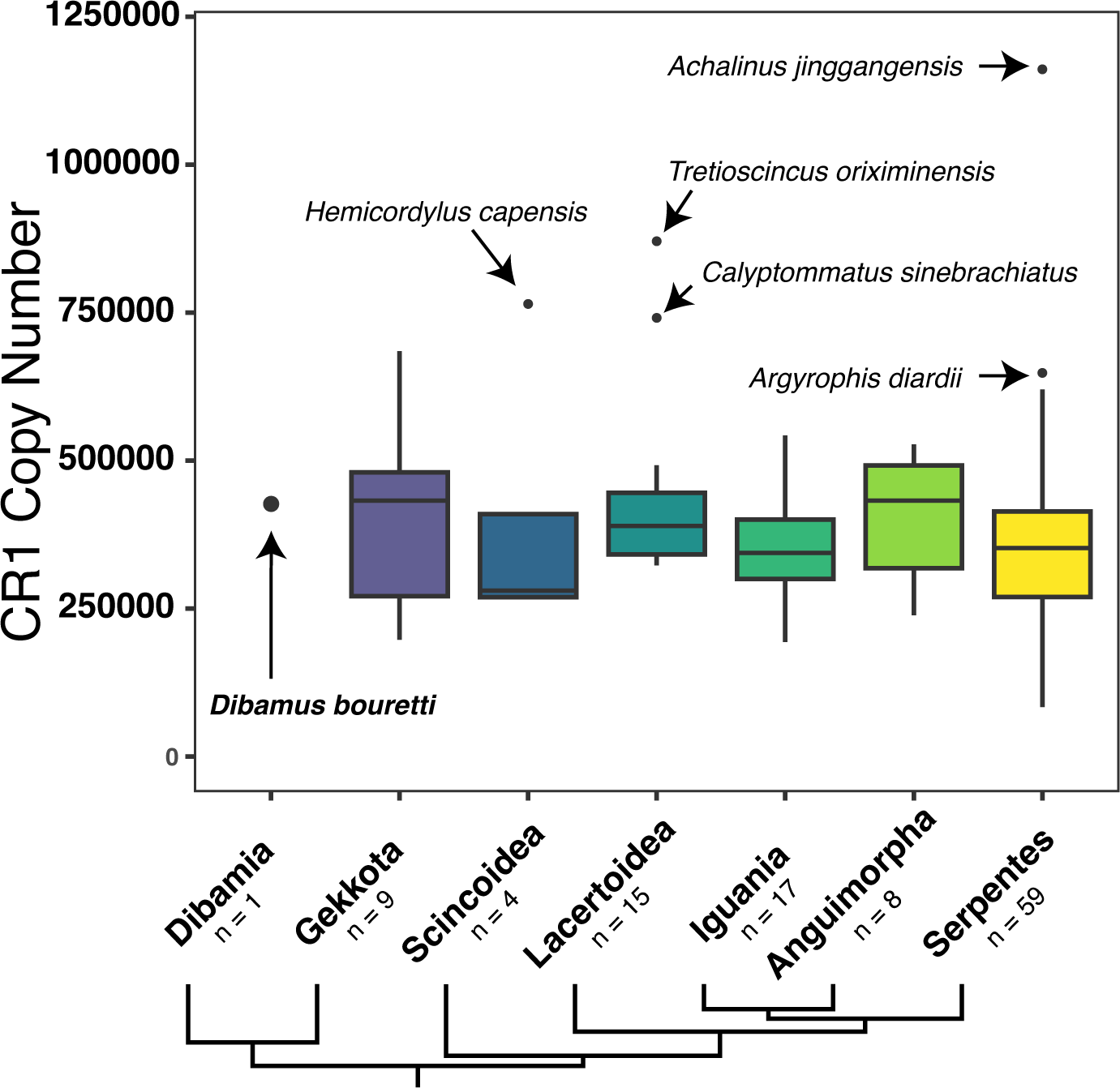
Variation in CR1 element copy number in 113 genomes from all 7 major squamate clades. The phylogenetic relationships and sample sizes for the seven major squamate clades (Dibamia, Gekkota, Scincoidea, Lacertoidea, Iguania, Angiumorpha, Serpentes) are given. Vertical bars represent median, small black circles represent outliers.

The amount of CR1 insertions in the genome varied dramatically between and within squamate clades (Fig. 1). Genomes of species classified in Lacertoidea (n=15) and Serpentes (n=59) showed the most variation in CR1 copy number with ranges of 547,627 inserts and 1,077,559 inserts, respectively. These large ranges are associated with the outliers in these groups, although Serpentes had a range of 536,785 even when excluding outliers. Serpentes included two outliers: *Achalinus jinggangensis* (family Xenodermidae; Zong and Ma 1983) with 1,161,213 CR1 copies accounting for over 18% of the genome, the most of any genome in our dataset, and the blind snake *Argyrophis diardii* (family Typhlopidae; Scolecophidia; Pyron and Wallach 2014) with 648,249 insertions. The outliers in Lacertoidea were the only two species from the lizard family Gymnophthalmidae, *Calyptommatus sinebrachiatus* and *Tretioscincus oriximinensis*. The outlier in Scincoidea, *Hemicordylus capensis*, is also the only representative of its family (Cordylidae), and therefore may not represent a true outlier due to the small sample size for Scincoidea (n=4).

CR1 copy number varied much more extensively across squamates compared to turtles and archosaurs (Supplementary Fig. 5). Both the maximum and minimum CR1 copy number for squamates were found in snakes, ranging from 1,161,213 in *Achalinus jinggangensis* to 83,654 in *Simalia boeleni* (family Pythonidae; Reynolds et al. 2014). The mean CR1 copy number in squamate genomes was 373,541.4, with a median of 361,430. In contrast, CR1 copy number in turtle genomes ranged 300,038 inserts, from 635,205 (*Pelodiscus sinensis*, family Tryonichidae, Cryptodira; Stuckas and Fritz 2011) to 335,167 inserts in *Dermatemys mawii* (family Dermatemydidiae, Cryptodira; González-Porter et al. 2013), with a mean of 482,047 and a median of 490,220. CR1 copy number in crocodilians showed the smallest range of 44,448 inserts, with a mean of 527,897.2 and median of 516,830.

CR1 copy number in avian genomes was the lowest across the sauropsid groups analyzed with a mean of 176,281 inserts and a median of 138,351, reflecting their small genome sizes and overall low TE abundance. However, there were five outlier bird species with much higher CR1 copy numbers, including the genomes of woodpeckers and barbets (Manthey et al. 2018). These outlier species contained an average CR1 copy number of 710,915.8, whereas average CR1 copy number in birds excluding outliers was 162,573. The maximum CR1 copy number in birds was in *Eubucco bourcierii* (Piciformes, family Capitonidae) with 896,033 inserts, over 200,000 inserts more than the maximum CR1 copy number in turtle genomes. The minimum CR1 copy number in birds was in *Parabuteo unicinctus* (Accipitriformes, subfamily Buteoninae) with 42,426 copies.

### Highly Heterogenous Rates of CR1 Copy Number Evolution in Squamates Compared to other Reptiles

We estimated the rate of CR1 copy number evolution in terms of number of inserts per million years for each branch of the species-level phylogenies for squamates, turtles, and archosaurs (including birds and crocodilians both together and separately) using ancestral state reconstruction (Fig. 2; Materials and Methods). The distributions of evolutionary rates for CR1 copy number from the squamate, archosaur, and turtle phylogenies, respectively, were all roughly bell-shaped around their means (Fig. 3); however, there were significant differences between all of them (p<0.005, D>0.25, nonparametric asymptotic 2-Sample Kolmogorov-Smirnov test).

**Figure 2:**
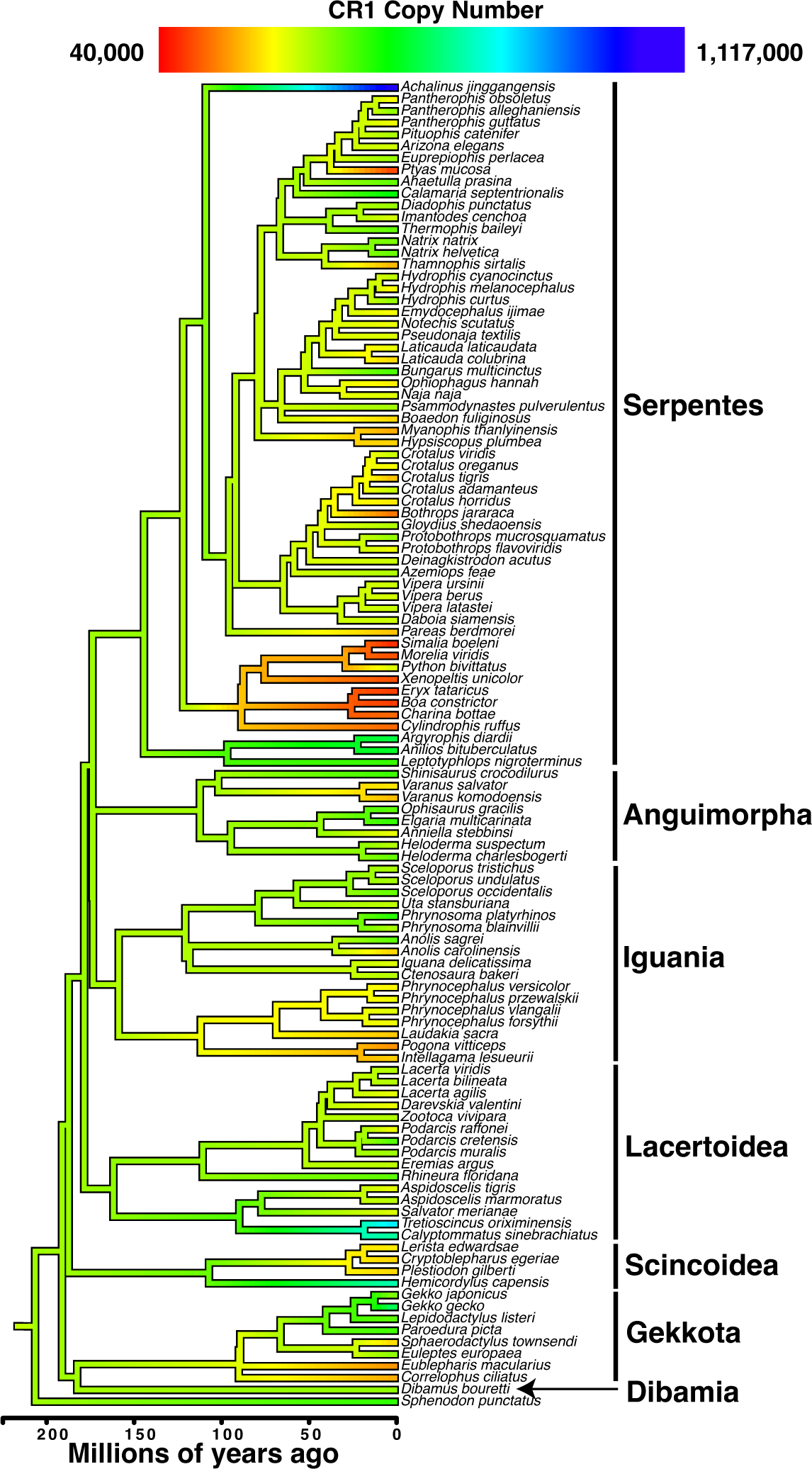
Species tree for 111 squamates representing seven major clades with ancestral state estimation for CR1 copy number mapped onto branches. Species tree was based on 2,999 nucleotide sequences of high-confidence single-copy orthologs (see Methods and Materials). The units for branch length are in terms of millions of years. Outgroups other than *Sphenodon* (*Homo sapiens, Gallus gallus, Alligator mississippiensis, Chrysemys picta*) not shown. CR1 insert number was modeled as a continuous trait for all genomes with annotated repeats using fastAnc in the package phytools (Revell 2012).

**Figure 3.**
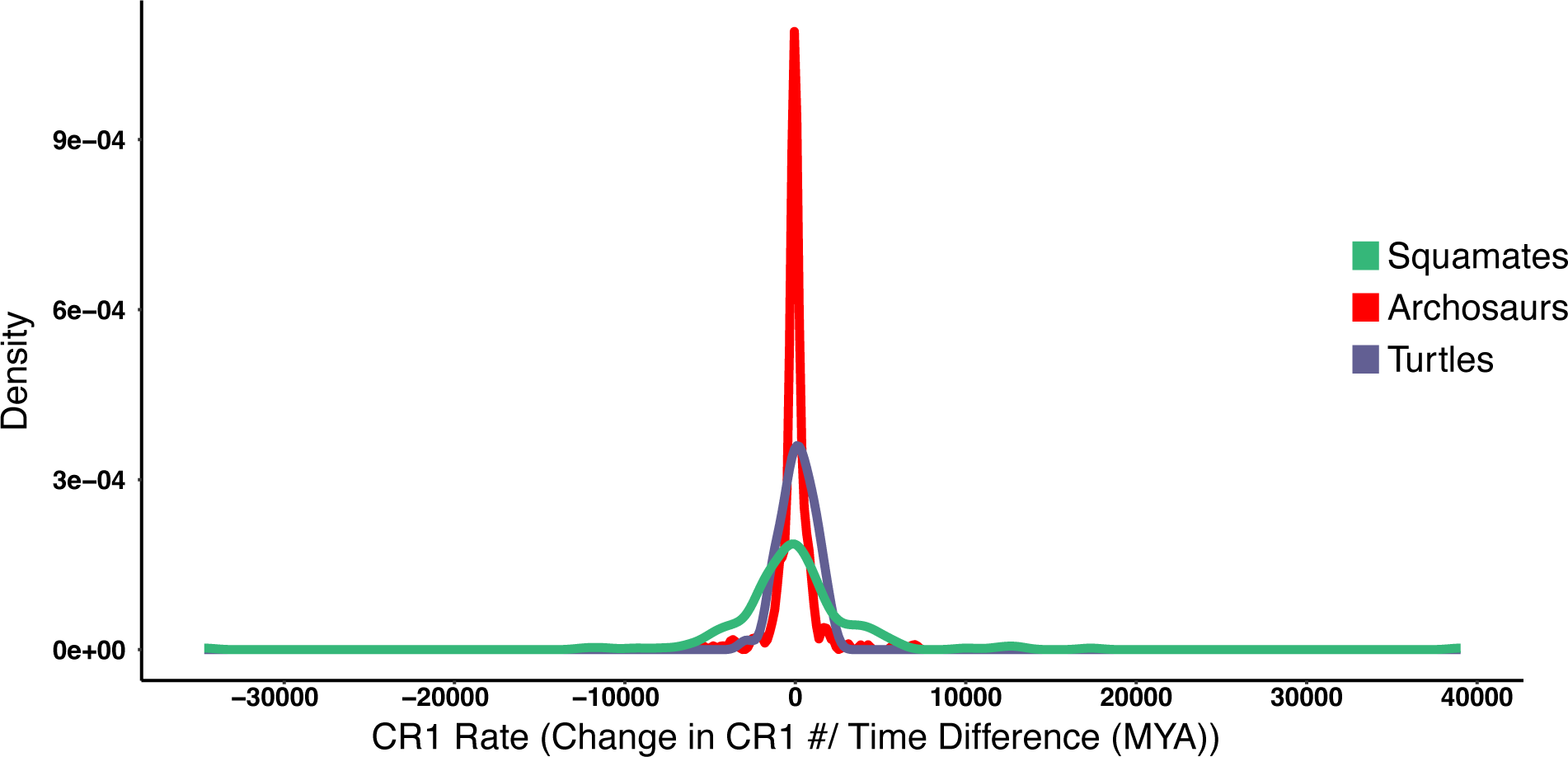
Estimated rates of CR1 copy number evolution are significantly different and more varied in squamates compared to other reptiles. Comparisons of these distributions were made using the Kolmogorov-Smirnov Test (P<0.0001). Overlaid density plot of normalized archosaur (purple), squamate (green), and turtle (blue) CR1 rates. Turtle CR1 rates range from −2,877.3 to 2,031.0 with a mean of 128.8 and a median of 146.8, while squamate CR1 rates range from −34,679.7 to 39,034.5 with a mean of −19.1 and a median of −156.2. Archosaur CR1 rates average −22.0 with a median of −36.4, and avian-only rates average −29.8 with a median of −38.6.

The estimated rates of CR1 copy number evolution for turtles were normally distributed (p=0.61, Shapiro-Wilk normality test) while estimated rates of CR1 copy number evolution for squamates, archosaurs, and the avian-only analysis were nonnormal (p<0.0001, Shapiro-Wilk normality test), consistent with residual analyses (Supplementary Fig. 7). The variances of each distribution were also heterogenous (p=0.0136, Levene’s test). Because the estimated rate of CR1 copy number evolution on a given branch was either positive (indicating overall gain from the ancestral state) or negative (indicating overall loss from the ancestral state), we alternatively log-transformed the data using log(1 + CR1 rate for the branch – the minimum CR1 rate). The Shapiro-Wilk normality test results for transformed data were: turtles, W = 0.36094, p = 1.362e-14; squamates, W = 0.11339, p < 0.001; archosaurs and avian-only, W = 0.23974, p < 0.001. The distribution of the estimated rates of CR1 copy number evolution for the squamate phylogeny was also significantly different from those of other reptiles based on transformed data (p < 0.0001, D= 0.99548, nonparametric asymptotic 2-Sample Kolmogorov-Smirnov test).

The estimated rate of CR1 copy number evolution for squamates was on average negative, ranging from −34,679.7 copies per million years (copies/MY) to 39,034.5 copies/MY, with a mean of −19.1 and a median of −156.2. Meanwhile, the estimated rate of CR1 copy number evolution for turtles was overall slightly positive, ranging from −2,877.3 copies/MY to 2,031.0 copies/MY, with a mean of 128.8 and a median of 146.8. Among archosaurs, the estimated rate of CR1 copy number evolution for birds was negative with a mean of −29.8 copies/MY and a median of −38.6; meanwhile, like turtles, crocodilians showed an average net increase in CR1 copy number with a mean of 276.9 copies/MY and a median of 417.6. Estimated rates of CR1 copy number evolution in archosaurs ranged from −6,062.4 copies/MY to 7,102.1 copies/MY, which was less than one-fifth the range for squamates.

### Episodic Bursts of CR1 Activity across Squamate Evolutionary History

In addition to overall CR1 copy number, as well as the estimated rate of CR1 copy number evolution being highly varied between squamates, we also found that CR1 activity was highly variable across squamate species and clades (Fig. 4). We used repeat landscapes of each analyzed genome to model CR1 activity over time, based on the Kimura 2-parameter (K2P) divergence of each CR1 insert from their family consensus sequence as a proxy for element age (see Materials and Methods). Among the repeat landscapes of CR1 divergence from representatives of ten squamate clades (Fig. 4), the scolecophidian snake *A. diardii* showed the largest proportion of the genome consisting of CR1 inserts at <10% divergence from consensus. In contrast, some squamate genomes contained relatively low amounts of CR1 insertions in more recent bins of K2P divergence. The *Boa constrictor* landscape revealed limited CR1 activity across all divergence bins, matching the pattern seen in other henophidian snake genomes which generally lack CR1 copies compared to other squamates (Supplementary Table 1). These results are consistent with the relative losses in CR1 copy number in snakes that we detected using ancestral state reconstruction (Fig. 2).

**Figure 4:**
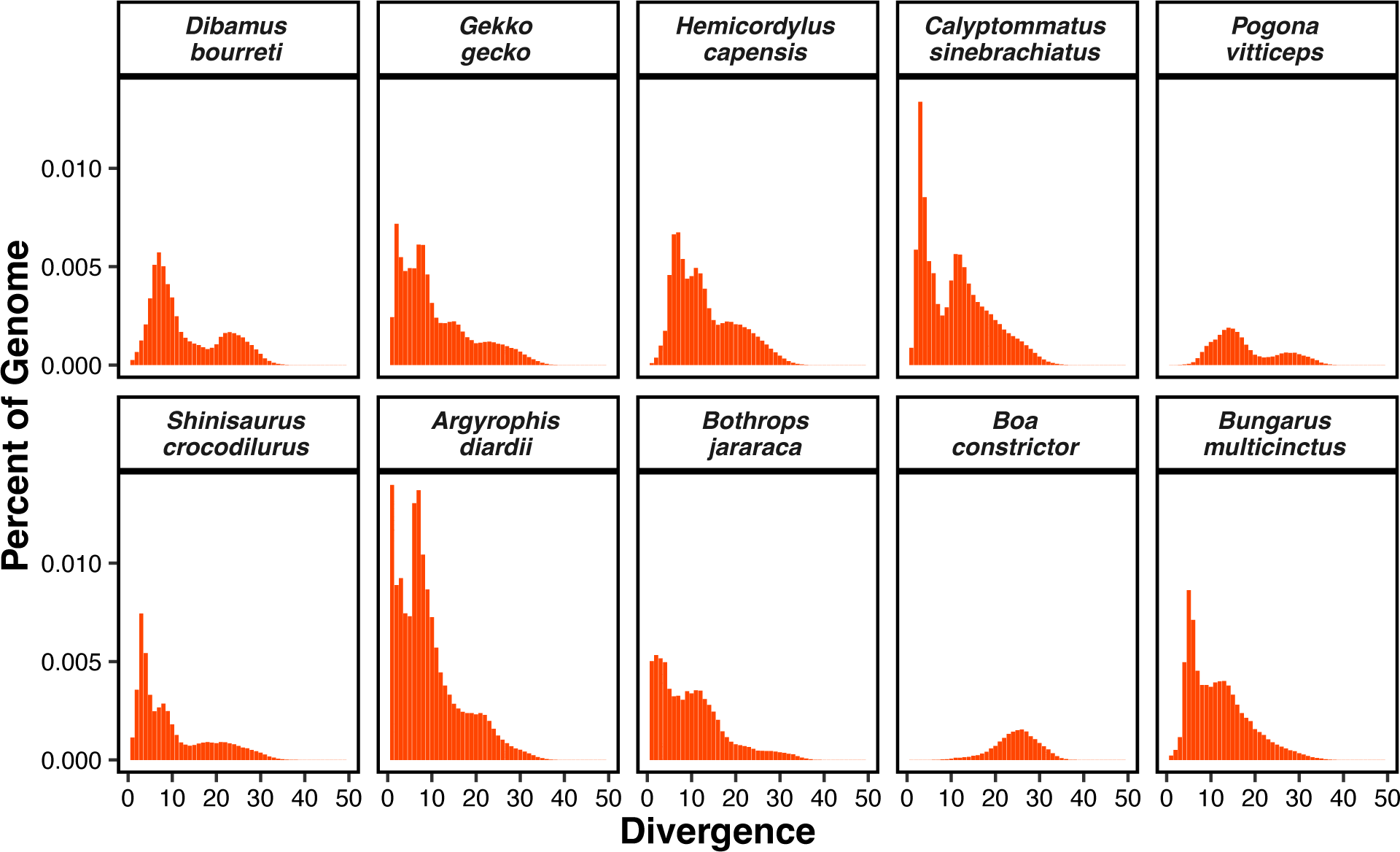
CR1 Repeat landscapes for 10 squamate species representative of major clades. Representatives were chosen primarily based on number of significant Wilcoxon pairwise comparisons (p<0.05) at 5% divergence, and secondarily based on assembly quality (e.g. N50, % gaps, completeness). Represented clades: Dibamia: *Dibamus bourreti*, Gekkota: *Gekko gecko*, Scincoidea: *Hemicordylus capensis*, Lacertoidea: *Calyptommatus sinebrachiatus*, Iguania: *Pogona vitticeps*, Anguimorpha: *Shinisaurus crocodilurus*; Serpentes [Scolecophidia: *Argyrophis diardii*, Viperidae: *Bothrops jararaca*, Henophidia: *Boa constrictor*, Elapidae: *Bungarus multicinctus*].

We used pairwise Wilcoxon rank-sum tests to compare all the distributions of CR1 K2P divergence calculated for 113 squamate genomes (Materials and Methods). The homalapsid snake *Myanophis thanlyinensis* (Köhler et al. 2021) had the most significant pairwise comparisons at 5% and 10% K2P divergence (i.e., ∼62.5 and ∼125 MYA, respectively, see Discussion), and the second most significant pairwise comparisons at 15% K2P divergence (Supplementary Table 1). Most species with the highest amount of significant pairwise comparisons were henophidian snakes (e.g., boas and pythons), reflecting the loss of CR1 copy number that is shared by these species (Fig. 2). CR1 landscapes of squamate species with some of the highest CR1 copy numbers included the xenodermid snake *Achalinus jinggangensis* and the gymnophthalmid lizards *Tretioscincus oriximinensis* and *Calyptommatus sinebrachiatus*, which were significantly different from most squamate landscapes across all divergence ranges. However, the cordylid *Hemicordylus capensis* (Scincoidea: Cordylidae), which contained a relatively higher number of insertions indicative of a large CR1 expansion, only had significantly different comparisons at the 10% and 15% levels of CR1 K2P divergence. This suggests some expansions of CR1 in cordylids may be older than in other squamates (∼187.5 MYA, see Discussion); however, more sampling from cordylids and other scincoids is required to more accurately determine the age of this expansion.

### CR1 Insertions Are Mostly Truncated and Are Found Far from Exons in Squamate Genomes

To understand how CR1 retrotransposons are distributed across squamate genomes, including their interactions with protein-coding genes, we analyzed the genomic locations of CR1 inserts, their lengths, and their proximity to gene (exonic) annotations in 30 squamate genome assemblies (Table 1; Supplementary Figure 10; see Materials and Methods). We found that in all squamate genomes the vast majority (97%) of CR1 inserts are truncated rather than full-length. The mean distance of CR1 inserts to exons in base pairs (49,144 bp) was greater than the mean intergenic distance overall in squamate genomes (41,330 bp) (Wilcoxon rank-sum test, z = −2.8909, p = 0.00386). The mean distance of CR1 inserts upstream of exons (52,227 bp) was greater than the mean distance of CR1 inserts downstream of exons (47,568 bp). We found a negative correlation between average distance in base pairs of CR1 inserts to exons and the average CR1 insert length in base pairs. In the analyzed squamate genomes we also found an average of 316 CR1 inserts which overlapped exons for a mean length of overlap of 239 bp. We did not detect a significant difference between average GC content 1 kb upstream or downstream of CR1 inserts (43%) and average total genomic GC content (42%) in the squamate genomes.

**Table 1.** CR1 insertion length, distance to exons, and overlap with exons data for 29 squamate genome assemblies.

| Major Clade | Family | Species | Mean CR1 Length (bp) | No. Full-Length CR1s | No. Truncated CR1s* | Mean CR1-Exon Distance (Absolute Value) (bp) | Mean Intergenic Distance (bp) | No. Overlapping CR1s-Exons |
| --- | --- | --- | --- | --- | --- | --- | --- | --- |
| Gekkota | Eublepharidae | *Eublepharis macularius* | 1,059 | 2,063 | 37,126 | 99,118 | 58,680 | 190 |
|  | Sphaerodactylidae | *Euleptes europaea* | 842 | 894 | 48,871 | 53,794 | 53,086 | 103 |
|  |  | *Sphaerodactylus townsendi* | 989 | 3,130 | 50,002 | 50,184 | 48,630 | 281 |
|  | Gekkonidae | *Gekko japonicus* | 903 | 1,805 | 67,648 | 48,602 | 43,394 | 153 |
| Scincoidea | Cordylidae | *Hemicordylus capensis* | 896 | 2,360 | 77,170 | 30,385 | 48,305 | 1,768 |
| Lacertoidea | Lacertidae | *Lacerta agilis* | 736 | 87 | 17,142 | 43,853 | 33,777 | 126 |
|  |  | *Lacerta bilineata* | 833 | 502 | 19,003 | 26,491 | 22,091 | 81 |
|  |  | *Lacerta viridis* | 805 | 367 | 20,723 | 36,909 | 25,969 | 65 |
|  |  | *Podarcis cretensis* | 929 | 1,367 | 24,939 | 41,934 | 35,101 | 208 |
|  |  | *Podarcis raffonei* | 992 | 1,743 | 24,141 | 39,256 | 35,028 | 260 |
| Serpentes | Colubridae | *Ahaetulla prasina* | 865 | 1,664 | 79,851 | 70,043 | 49,209 | 744 |
|  |  | *Pantherophis alleghaniensis* | 730 | 57 | 36,399 | 40,117 | 37,592 | 256 |
|  |  | *Pantherophis guttatus* | 736 | 79 | 32,585 | 38,855 | 36,644 | 216 |
|  |  | *Pantherophis obsoletus* | 729 | 0 | 32,775 | 40,052 | 37,091 | 181 |
|  |  | *Pituophis catenifer* | 762 | 244 | 36,440 | 58,946 | 48,621 | 402 |
|  | Elapidae | *Notechis scutatus* | 1,066 | 3,797 | 53,031 | 53,753 | 37,403 | 252 |
|  |  | *Pseudonaja textilis* | 953 | 2,181 | 48,730 | 54,312 | 39,363 | 293 |
|  | Pythonidae | *Python bivittatus* | 758 | 15 | 13,589 | 18,901 | 17,756 | 143 |
|  | Natricidae | *Thamnophis elegans* | 703 | 322 | 48,938 | 63,500 | 48,155 | 303 |
|  | Viperidae | *Crotalus adamanteus* | 977 | 2,788 | 62,564 | 22,951 | 23,369 | 477 |
|  |  | *Crotalus oreganus* | 906 | 1,326 | 48,528 | 51,874 | 42,640 | 405 |
|  |  | *Crotalus tigris* | 977 | 1,973 | 44,322 | 39,623 | 31,008 | 363 |
|  |  | *Gloydius shedaoensis* | 919 | 1,911 | 66,821 | 48,825 | 37,989 | 439 |
|  |  | *Protobothrops mucrosquamatus* | 809 | 530 | 52,118 | 34,806 | 27,115 | 91 |
| Anguimorpha | Helodermatidae | *Heloderma suspectum* | 1,153 | 5,960 | 73,910 | 61,733 | 84,786 | 115 |
|  | Varanidae | *Varanus komodoensis* | 919 | 1,474 | 56,387 | 52,828 | 41,144 | 476 |
|  |  | *Varanus salvator* | 1,082 | 4,329 | 56,221 | 78,707 | 52,414 | 432 |
| Iguania | Dactyloidae | *Anolis carolinensis* | 1,010 | 1,756 | 31,640 | 33,444 | 45,471 | 434 |
|  | Agamidae | *Laudakia sacra* | 1,351 | 6,516 | 40,026 | 80,867 | 61,179 | 153 |
|  |  | *Pogona vitticeps* | 802 | 54 | 18,032 | 59,646 | 36,883 | 71 |
| Average: |  |  | 906 | 1,710 | 43,989 | 49,144 | 41,330 | 316 |

\*Only CR1 inserts >500bp were considered.

### Conservation And Loss of CR1 Subfamilies across Reptile Evolution

In order to model the evolutionary history of full-length, active CR1 sequences that were producing new copies throughout the history of amniotes, we extracted, aligned, and filtered CR1 ORF2 reverse transcriptase sequences from each squamate genome, and used maximum-likelihood phylogenetic inference to cluster the sequences into subfamilies. We incorporated consensus sequences of 118 CR1 subfamilies identified in lepidosaurs with 106 additional CR1 consensus sequences from amniotes and an outgroup amphibian (*Xenopus tropicalis*) from RepBase (Bao et al. 2015) also using maximum-likelihood phylogenetic reconstruction (Supplementary Table 1). CR1 subfamily consensus sequences from RepBase consisted of 36 avian subfamilies, 69 crocodilian subfamilies, 15 turtle subfamilies, and 3 mammal subfamilies.

Our phylogenetic analysis revealed evidence of multiple lineages of CR1 inhabiting the genomes of amniotes (Fig. 5). Many CR1 subfamilies recovered from crocodilian, mammalian, turtle, and lepidosaurian genomes clustered together in a pattern that was inconsistent with the host phylogenetic relationships. Crocodilian CR1 subfamilies were found in numerous places along the amniote CR1 phylogeny, including 60 monophyletic CR1 subfamilies we recovered only from crocodilian genomes (Fig. 5, light purple). CR1 consensus sequences from mammalian and turtle genomes were scattered across the CR1 phylogeny, while our analysis of the tuatara genome revealed several tuatara-specific CR1 subfamilies that clustered with CR1 sequences from mammalian, crocodilian, and turtle genomes. All avian CR1 subfamilies were grouped together (Fig. 5, light red).

**Figure 5:**
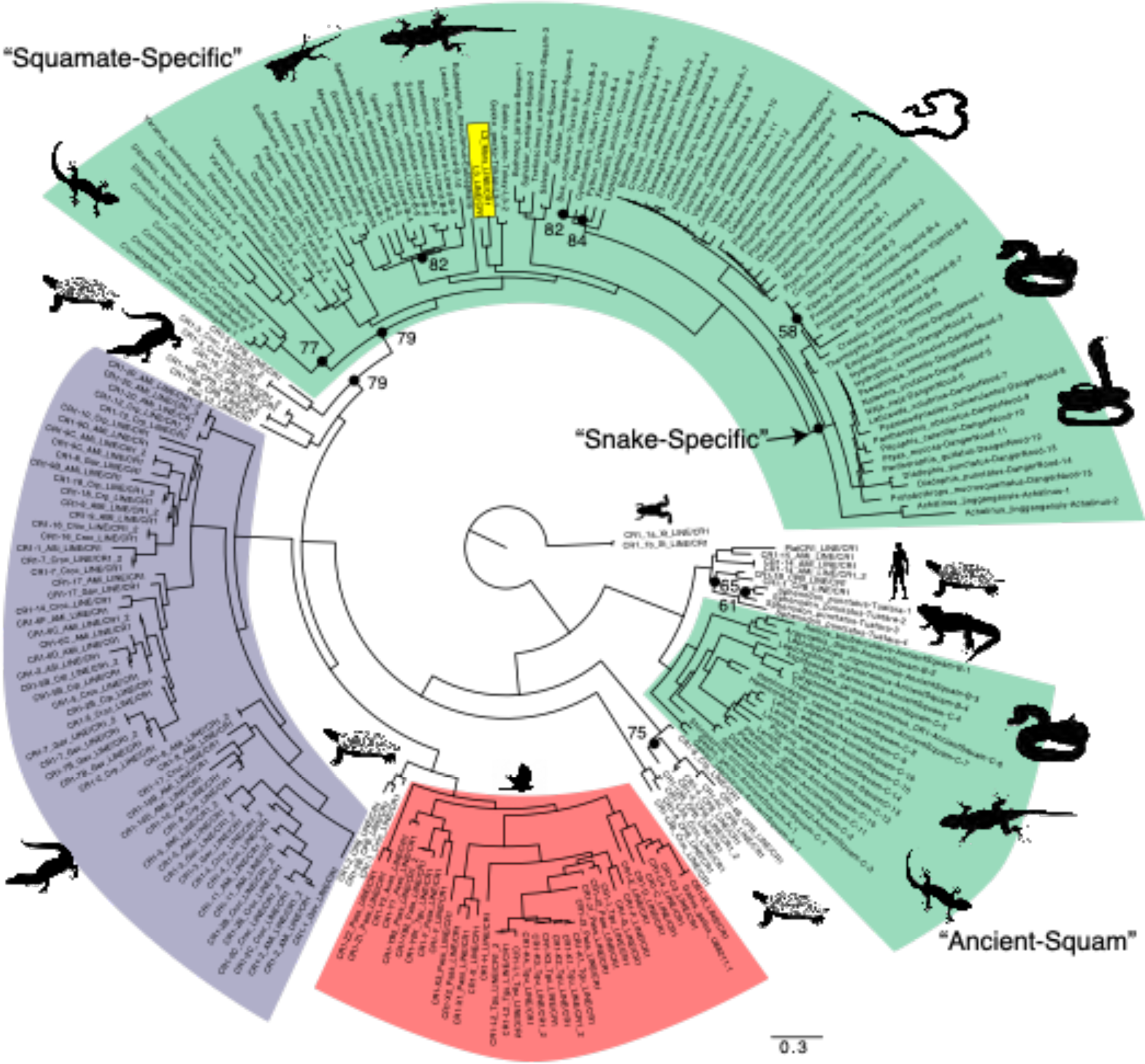
Maximum-likelihood phylogeny of 244 amniote CR1 subfamily consensus sequences with new squamate CR1 subfamilies labeled. Branch lengths are in terms of substitutions per site. Squamate subfamilies (light green) were named based on the clade most inclusive of the species represented in each CR1 subfamily. We identified 114 total squamate CR1 subfamilies from 63 squamate species, along with 4 total tuatara CR1 subfamilies. There were 36 total avian subfamilies (light red), 69 crocodilian subfamilies (light purple), 15 turtle subfamilies (not colored), and 3 mammal subfamilies (not colored). Within the Squamate-Specific clade there were L3 sequences, a subtype of CR1 found in mammals (Gentles et al. 2007), that clustered with a consensus sequences from a gecko (yellow box). Bootstrap values <90% are shown. RepBase abbreviations: “Plat” = Platypus, “Ami” = *Alligator mississippiensis*, “CPB” = *Chrysemys picta bellii*, “Croc” = Crocodilian, “Crp” = *Crocodylus porosus*. Animal silhouettes from www.phylopic.org under the public domain.

Our analysis enabled a deeper look into CR1 evolution in squamates that has not been possible until recently. The 114 CR1 subfamilies we identified in squamates could be classified into two main clades (Fig. 5, light green). The first is an ancient CR1 clade (“Ancient-Squam”, 100% bootstrap support) comprised of consensus sequences we collected from 16 squamate genomes that forms the sister taxon to the clade of CR1 subfamilies collected from the tuatara, turtle, crocodilian, and mammalian genomes, which together formed an outgroup to all other amniote CR1 consensus sequences. The Ancient-Squam CR1 clade contained consensus sequences recovered from at least one species from each of the seven major squamate clades (Iguania, Anguimorpha, Serpentes, Gekkota, Dibamia, Lacertoidea, Scincoidea).

The remaining CR1 consensus sequences from squamate genomes formed a large clade of CR1 subfamilies that was unique to squamates, including sequences recovered from species across the squamate phylogeny (“Squamate-Specific”, 79% bootstrap support). Within this group, several CR1 subfamilies found in squamates clustered in patterns that were consistent with host phylogenetic relationships (see Discussion). These included CR1 subfamilies Viperid-A and Viperid-B that were unique to viperid snakes. Lizard-A sequences found in the dibamid (*Dibamus bouretti*) genome clustered with CR1 retrieved from several gecko genomes, which formed the outgroup to all other squamate-specific CR1 subfamilies.

There were several areas of the tree showing high-support grouping of subfamilies from species with large phylogenetic distances (e.g., Squam, Toxico-A, Toxico-B). While CR1 subfamilies from the Ancient-Squam clades were found only in squamate genomes, the Squamate-Specific clade contained one exception: an L3 subfamily from a mammalian host (L3_Mars_LINE/CR1 and L3_LINE/CR1) that was sister to the Tokay-L3 subfamily found in a gecko genome (*Gekko gecko*) with 100% branch support (Fig. 5, yellow box).

## Discussion

Our study has revealed contrasting patterns of diversification, activity, conservation, and loss of the CR1 retrotransposon family throughout amniote and specifically squamate evolution. We found that (1) CR1 copy number varies significantly across the genomes of squamates, and that this variation is much more extensive than in other reptile groups; (2) there has also been much more extensive variation in CR1 rates of evolution and CR1 retrotransposition activity across squamates compared to other reptiles; and (3) squamate genomes contain a diversity of shared and derived CR1 subfamilies indicative of both the persistence and evolutionary loss of several ancient CR1 lineages over amniote evolution. We also found that CR1 elements are mostly truncated in sequence and tend to be far upstream or downstream of exons in squamate genomes. These results point to a dynamic genomic environment in squamates that stands out in comparison to other sauropsids, that may shed light on the evolutionary forces that govern genome size and structure in vertebrates.

Comparative genomics studies which incorporate a large amount of publicly available data are subject to variability in data quality. We addressed these limitations in numerous ways. First, we filtered our whole genome assembly dataset for contiguity (i.e., N50>15 kbp) and predicted gene content (i.e., percent of complete and single-copy BUSCOs > 40%), which are associated with assembly quality (Thrash et al. 2020), except where we prioritized taxonomic representation (e.g., *Dibamus, Anilios, Myanophis*). Also, all comparative methods rely on accurate species trees, so for species tree reconstruction we extensively filtered orthologous gene alignments for length, gaps, and taxonomic coverage (see Materials and Methods). For CR1 phylogenetic analysis, we targeted only full-length ORF2 regions with verified presence of a reverse transcriptase and endonuclease region. To maximize the observable genomic changes along branches of the amniote phylogeny, we prioritized broad taxonomic representation. We suggest that future studies consider the tradeoffs inherent to achieving a balance between data quantity for statistical power and data quality for reproducibility. In particular, the greater use of long-read sequencing technologies will improve TE annotation going forward (Gable et al. 2023; Sproul et al. 2023).

Throughout >300 million years of amniote evolution, the number of CR1 retrotransposon copies inhabiting and diversifying within amniote genomes has varied widely, reflecting different modes of genome evolution among mammals, birds, and reptiles. Our estimated rates of CR1 evolution support a “slow and steady” mode of genomic evolution for turtles and crocodilians, consistent with molecular clock-based analyses based on fourfold degenerate sites (Shaffer et al. 2013; Green et al. 2014; Tollis et al. 2017; Gemmell et al. 2020). Meanwhile, avian genomic evolution, which parallels squamate evolution in terms of species diversity (i.e., both clades have ∼11,000 extant species), features a much smaller range of CR1 copy numbers than squamates, with few outliers (Manthey et al. 2018, Galbraith et al. 2021). The much wider variation in CR1 copy number, and the distribution of CR1 evolutionary rates, in squamates suggests many rapid shifts in genome evolution in this group of reptiles.

Most of the squamate genomes that we analyzed contained a relatively high proportion of CR1 insertions between 0-10% K2P divergence from consensus (Fig. 3; Supplementary Fig. 4), suggesting relatively recent CR1 activity. Applying a conservative genome-wide substitution rate of 0.0008 substitutions per site per million years for squamates (Perry et al. 2018; Finger et al. 2022), we estimate this range of CR1 divergence reflects CR1 activity within the last 125 million years of squamate evolution, which is younger than the divergence of the seven major squamate clades analyzed here (Fig. 2; Gamble et al. 2015; Pyron 2017; Burbrink et al. 2020; Simões et al. 2020; Brownstein et al. 2023). These results suggest that differences in the success of CR1 retrotransposons across squamates may be due to lineage-specific differences in CR1 activity over relatively short timespans.

While our analyses support the hypothesis that the common ancestor of amniotes hosted multiple subfamilies of CR1 in its genome, followed by multiple losses during amniote evolution (Suh et al. 2015), we have expanded into squamate taxonomic diversity to reveal a more dynamic CR1 evolutionary history than has been previously suggested. For instance, the Ancient-Squam CR1 clade contained consensus sequences from at least one species from each of seven major squamate clades (Iguania, Anguimorpha. Serpentes, Gekkota, Dibamia, Lacertoidea, Scincoidea), suggesting that this CR1 subfamily originated in the genome of the common ancestor of all extant squamates and has persisted for as much as 200 million years (Brownstein et al. 2023). This conservation of the Ancient-Squam CR1 clade in the genomes of some species is contrasted by its apparent evolutionary loss in the genomes of squamates where we did not find Ancient-Squam CR1 sequences.

Certain CR1 subfamilies in the Squamate-Specific clade phylogenetically clustered in patterns that are reminiscent of the host phylogeny, supporting a model of CR1 vertical transmission between ancestor to descendent genomes. For instance, Lizard-A sequences found in the dibamid (*Dibamus bouretti*) genome clustered with CR1 subfamilies retrieved from several gecko genomes, which in our analyses formed the outgroup to all other squamate-specific CR1 subfamilies (with 79% bootstrap support). This result mirrors most species trees for squamates published in the molecular era, which place geckos and/or dibamids at the root of the squamate phylogeny (Pyron et al. 2013; Streicher and Wiens 2017; Burbrink et al. 2020; Singhal et al. 2021). We also recovered a large snake-specific CR1 clade with relatively short branch lengths throughout, which possibly coincided with the snake radiation that started ∼150 million years ago (Fig. 2, Caldwell et al. 2015). The shortest branch lengths in the squamate CR1 phylogeny were all in venomous snake CR1 subfamilies, perhaps reflecting the rapid diversification of those snake families during the late Cretaceous and Paleogene Periods (Pyron and Burbrink 2012; Lee et al. 2016). In contrast, some species of squamates contained groups of CR1 subfamilies that were reciprocally monophyletic (i.e., found only in that species), such as *Correlophus ciliatus*, *Dibamus bouretti*, and *Anolis carolinensis*, indicating more recent diversification of CR1 subfamilies unique to these species’ evolutionary histories.

Our reconstruction of CR1 evolution relies on the assumption that CR1 is vertically transmitted between ancestor and descendant genomes through the germline, rather than horizontally transferred between very distantly related host taxa. Since phylogenetic reconstructions of non-LTR retrotransposons generally follow their host phylogenies (Malik et al. 1999; Kordiš et al. 2006; Waters et al. 2007; Boissinot and Sookdeo 2016), this model of vertical transmission (with occasional discordance due to incomplete lineage sorting, introgression, and loss via deletion) is reasonable most of the time. However, horizontal transfer of non-LTR retrotransposons has been demonstrated in reptiles, as in the transfer of Bov-B elements between boid snakes and a ruminant mammalian ancestor (Ivancevic et al. 2018), as well as between squamates and potentially several ectoparasitic hosts (Pasquesi et al. 2018). In particular, horizontal transfer of CR1 has occurred between *Maculinea* butterflies and *Bombyx mori* silk moths (Novikova et al. 2007). In general, these instances of horizontal transfer can be detected when the element phylogeny is inconsistent with the host species phylogeny. In our phylogenetic analysis of CR1 consensus sequences, the CR1/L3 subfamily identified in the Tokay gecko (*Gekko gecko)* clustered with mammalian L3 deep within the otherwise squamate-specific CR1 clade (Fig. 5). Previously, it was suggested that this pattern of a mammalian L3 grouping with lepidosaurian CR1 sequences supported a model of shared ancestry followed by loss in mammals and other amniotes (Suh et al. 2015). After expanding the taxonomic sampling of squamates for our study, we suggest a more parsimonious explanation may be horizontal transfer between a squamate and a mammalian host.

Variation in dynamics of TE movement has been identified as a main factor in determining eukaryotic genome size (Gregory 2002; Blommaert 2020). Comparative research in birds and mammals support an “accordion” model whereby DNA loss counteracts TE expansion (via transposition) and results in stable genome sizes over evolutionary timescales. Across squamates, genome size is tightly constrained (Pasquesi et al. 2018; Armstrong et al. 2020; Gable et al. 2023), showing little variation across large phylogenetic distances. Our results suggest that while CR1 has been highly active over squamate evolutionary history, there is also evidence for the loss of CR1 subfamilies in the genomes of some squamate lineages, supporting an accordion model for squamates. For instance, we estimated that the net rate of CR1 copy number evolution for the entire squamate phylogeny is negative. However, not much is known about patterns of DNA deletion in squamates. The role of deletions in maintaining genome size, whether by non-allelic recombination resulting from the accumulation of TEs such as CR1, or via smaller deletions at the base-pair level, will shed light on the mechanisms that control genome size variation in amniotes.

CR1 has been highly active in squamates and has produced large copy numbers in squamate genomes; however, similar to initial studies of the green anole genome (Novick et al. 2009; Alföldi et al. 2011) we found that on average only ∼3% of CR1 inserts were full-length (and therefore potentially active) across squamate genomes. This may be due to: (1) the retrotransposition process being mistake-prone and failing to produce many full-length CR1 insertions; (2) shorter elements being the result of DNA deletions; or (3) selection which constrains the number of full-length elements in the genome. While the other two choices are possibilities, we think purifying selection (the third choice) plays a large role in squamate TE dynamics. In natural populations of green anoles, full-length non-LTR retrotransposon insertions are found at significantly lower allele frequencies than truncated ones (Tollis and Boissinot 2013; Ruggiero et al. 2017; Bourgeois et al. 2020), suggesting selection acts against TEs according to their length, possibly due to the deleterious effects of ectopic recombination (Petrov et al. 2003; Boissinot et al. 2006). We found that across squamate genomes CR1 insertions were relatively far from exons, suggesting selection against TE insertions in genic regions. Truncated CR1 elements may be slightly more tolerated near genes, as well as genome-wide, due to weaker purifying selection, perhaps because they are more likely to become fixed in natural populations of lizards (Tollis and Boissinot 2013; Ruggiero et al. 2017; Bourgeois et al. 2020). This would explain why CR1 elements persist in squamate genomes despite purifying selection widely acting against TE accumulation.

The mutational effect of TE activity can be deleterious but also sometimes beneficial to the host genome, resulting in new phenotypes and positive selection (Pastuzyn et al. 2018; Rishishwar et al. 2018; Schrader and Schmitz 2019). We found that CR1 insertions were much closer to genes on average in the genome of the scincoid *Hemicordylus*, which also contained the highest number of CR1 insertions overlapping exons (1,768). Understanding the contribution of TEs to exon sequences will be important for investigations of the evolution and genetic regulation underlying fascinating squamate phenotypes, including leglessness (Roscito et al. 2022). The variation in squamate CR1 evolution uncovered in this study may also explain the unmatched abundance of microsatellites in squamate genomes (Adams et al. 2016), particularly in snakes (Pasquesi et al. 2018) where CR1 elements often contain microsatellites on their 3’ ends (Castoe et al. 2011). Understanding the role of CR1 activity in microsatellite exaptation, and protein-coding and regulatory regions in general, will shed light on CR1’s potential to be a mechanism of phenotypic evolution in squamates and other vertebrates.

TE evolution directly impacts genome architecture and drives genetic diversity across eukaryotes. Using expanded genomic datasets for reptiles, we uncovered evidence of evolutionary gain and loss of CR1 retrotransposon diversity throughout amniote genome evolution, identifying unique patterns of CR1 diversity and activity in squamate genomes. With diverse TE profiles and evidence of recent TE activity, the genomic ecosystem of squamates appears less constrained than all other amniotes. Our study underscores squamates as a promising research model for not only TE activity and diversity, but also outstanding questions in the evolutionary dynamics of genome size, structure, and function in vertebrates.

## Materials and Methods

### Genomic Data

We downloaded whole genome assemblies for 113 squamate species and the tuatara (*Sphenodon punctatus*, Accession: GCA_003113815.1) from NCBI (Sayers et al. 2022), Genome Warehouse (Chen et al. 2021), DNA Zoo (Dudchenko et al. 2017), and other open-source data repositories from individual publications (see Supplementary Table 1). For species with multiple assemblies available, the representative genome was chosen based on highest contig N50 (Bradnam et al. 2013). In total, we collected 114 lepidosaur genomes. We also downloaded whole genome assemblies for 30 turtle, 200 avian, and 4 crocodilian species from NCBI (see Supplementary Table 1).

### De Novo Repeat Annotation

We identified repetitive elements in each genome assembly using the *de novo* repeat finder RepeatModeler2.0 (Flynn et al. 2020) to build species-specific repeat libraries with search engine option “ncbi”. We then used RepeatMasker4.1 (Smit et al. 2013) to query each species-specific repeat library against the appropriate reference genome, generating repeat annotations for each assembly using the options “-xsmall”, “-no_is”, and “-a” to generate a *.align alignment output file for downstream analyses. To remove nested elements from TE annotations, a Perl script from Kapusta et al. 2017 (parseRM.pl, available at https://github.com/4ureliek/Parsing-RepeatMasker-Outputs) was used to parse *.align files. We calculated Kimura 2-parameter (K2P) corrected percentages of divergence from TE family consensus sequences, using calcDivergince.pl in RepeatMasker, accounting for CpG sites. All CR1 landscapes were generated using a custom R script. We used pairwise Wilcoxon rank-sum tests in R to compare the CR1 landscapes of 113 squamate species. We focused on the total proportion of the genome consisting of CR1 insertions in bins of 5%, 10%, and 15% K2P divergence to capture recent activity, and used a false-discovery rate p-value adjustment for multiple comparisons (p.adjust.method = “BH”) (Benjamini and Hochberg 1995).

### Squamate, Turtle & Archosaur Species Tree Inference

To infer ancestral states across reptiles, we generated a species tree using markers from whole genome sequence for 111 squamates, with the tuatara, the painted turtle (*Chrysemys picta*), the American alligator (*Alligator mississippiensis*), the chicken (*Gallus gallus*), and human for outgroups (Supplementary Table 1). Two squamate genomes were excluded from species tree inference due to low genome completeness where high-quality taxonomic representatives of those lineages were available (*Gonatodes ferrugineus* and *Thamnophis elegans*). We extracted nucleotide sequences of 7,453 high-confidence single-copy sauropsid orthologs using BUSCO v 5.4.4 (OrthoDB v10, Manni et al., 2021). We used MAFFT v7.505 (Katoh & Standley, 2013) to generate alignments for each ortholog, and TrimAL v1.4.rev15 (Capella-Gutierrez et al., 2009) to impose a gap threshold set to remove all positions in alignments with gaps in 30% or more of sequences (-gt 0.7). We generated summary statistics for all alignments with AMAS v3.04 (Borowiec, 2016), and filtered for alignments with less than 15% gaps, greater than 1,500 bp in length, less than 15,000 bp in length, and a minimum of 80% taxa representation.

After filtering, we analyzed 2,999 orthologs totaling 8,625,493 sites in IQ-Tree v2.2.0.4 (Minh et al., 2020) to infer gene trees using maximum likelihood with ModelFinder implemented to determine the best-fit model for each partition (see Supplementary Materials). The resulting gene trees were used to infer a coalescent-consistent multilocus species tree in ASTRAL-III (C. Zhang et al., 2018), with branch support measured in local posterior probabilities (Sayyari and Mirarab 2016). The species tree was rooted using the *H. sapiens* outgroup. All nodes were fully supported based on local posterior probabilities, aside from three nodes known to be areas of phylogenetic conflict within squamates (Supplementary Fig. 1): Dibamia-Gekkota (66%, Burbrink et al. 2020; Singhal et al. 2021), the Iguania-Anguimorpha (91%, Losos et al. 2012; Simões et al. 2018), and the Pseudaspididae-Elapidae (43%, Zaher et al. 2019).

The turtle coalescent-based phylogeny was inferred with the same methods described above, with genomes of 30 turtle species and five outgroups (the tuatara, the green anole (*Anolis carolenensis*), the American alligator (*Alligator mississippiensis*), the chicken (*Gallus gallus*), and human; Supplementary Table 1). We extracted nucleotide sequences of 7,448 high-confidence single-copy sauropsid orthologs, and after filtering trimmed alignments based on criteria described above, a total of 3,342 orthologs with 9,716,540 sites were used to infer gene trees and the resulting coalescent-based species tree (rooted with *H. sapiens*). The coalescent-based species tree for turtles fully recovered the Lepidosauria, Archelosauria, Pleurodira, and Cryptodira clades (Shaffer et al. 2017; Gable et al. 2022). All nodes were fully supported based on local posterior probabilities (Supplementary Fig. 2).

We inferred the archosaur coalescent-based phylogeny with the same methods described above, using 200 bird species and four crocodilian species, as well as four outgroups (the tuatara, the green anole, the painted turtle, and human). We extracted nucleotide sequences of 8,311 high-confidence single-copy orthologs using BUSCO v 5.4.4 (OrthoDB v10 avian database). We used a total of 2,981 filtered orthologs with 8,417,077 sites. Most branches in the archosaur tree received full branch support, apart from order-level relationships in Neoavian birds (Supplementary Fig. 3), which are known areas of uncertainty in avian phylogenetics (Prum et al. 2015; Kuhl et al. 2021). The few areas of low branch support did not impact the estimates of CR1 evolutionary rate variation, given that bird genomes largely lack variation in repeat content (Kapusta and Suh 2017, Supplementary Table 1).

### Divergence Time Estimation

We estimated divergence times based on the coalescent-based squamate, turtle and archosaur phylogenies separately using the penalized-likelihood approach implemented in treePL (Smith and O’Meara 2012). We used TreePL with minimum and maximum fossil calibrations for five nodes (Amniota: minimum 318 millions of years ago or My, maximum 332.9 My; Sauropsida: minimum 255.9 My, maximum 295.9 My; Archosauria: minimum 247.1 My, maximum 260.2 My; Lepidosauria: minimum 238 My, maximum 252.7 My; Squamata: minimum 168.9 My, maximum 209.5 My) from Benton et al. 2015. An optimal smoothing parameter was chosen based on cross-validation testing, which sequentially removes terminal taxa to produce an estimate of branches from remaining data given an optimal smoothing parameter, which is found from the fit of the true branch and the pruned branch (Sanderson 2002). The *randomcv* option was used to run randomly sampled cross validation analysis, and the *prime* option was initially run to find the best values for *opt*, *optad*, and *optcvad* optimizers. We also used the *thorough* option to ensure the run iterated until convergence.

### Ancestral State Estimation Using CR1 Copy Numbers

To investigate differences in reptile CR1 evolution, we modeled CR1 copy number as a continuous trait to trace the evolution of this element across reptile diversification. We calculated CR1 copy number for each species using RepeatMasker annotation files to generate a data frame of trait values and used our species tree to inform taxa relationships. Any duplicate CR1 sequences in repeat annotations were removed using seqkit “rmdup” command. We first inferred ancestral states of CR1 insert copy numbers on each branch of the species-level phylogenies for squamates, turtles, and archosaurs (birds and crocodilians both together and separately) with ancestral state reconstruction using fastAnc in the phytools R package (Revell, 2012) with the species trees and CR1 trait file as input.

Within each phylogeny, we estimated the rate of CR1 copy number evolution for each branch by calculating the estimated CR1 copy number on the younger node minus the estimated CR1 copy number on the older node, divided by the length of the branch in millions of years (i.e., Δ CR1 copy number/ Δ Time (My); Supplementary Materials). Because the fastAnc method assumes homogenous rates across a phylogeny, we tested for evidence of rate heterogeneity in CR1 evolution using the Savage-Dickey metric implemented in the evorates R package (Martin et al. 2023) and found no significant evidence of heterogeneity. We set variance and confidence intervals to ‘TRUE’ (i.e., vars=TRUE, CI=TRUE) to measure variability in estimates.

### Phylogenomic Estimation of Squamate CR1 – Within-Species Analyses

To reveal the phylogenetic structure of CR1 elements found in squamate genomes, we inferred CR1 phylogenies using full-length, active CR1 elements from the genomes of 78 squamate species representing 25 families and all 7 major clades (Supplementary Table 1; Burbrink et al. 2020; Singhal et al. 2021). Our full dataset of 111 squamate genomes was filtered for taxonomic representation primarily and assembly contiguity (i.e., N50) secondarily to reduce computational burden in manual curation of consensus sequences. In order to extract CR1 sequences from each assembly, we first converted the RepeatMasker *.out output files to standard BED annotation files via the ‘rmsk2bed’ command in ‘bedops’ (Neph et al. 2012) with option “—keep-header”. Family-specific BED files of full-length CR1 elements were generated using bash scripting, with a minimum cut-off value of 2000 bp used to initially filter for at least 60% complete ORF2 region (see Supplementary Table 1 for cut-offs used for each assembly) and maximum 6000 bp cut-off to remove artifactual sequencing errors (e.g., chimeric sequences). The CR1 ORF2 is known to be highly conserved across species due to the presence of an RT (reverse transcriptase) region, so the ORF2 is commonly isolated for comparative studies (Novick et al. 2009). We then extracted CR1 sequences for each species using *bedtools getfasta* with bedtoolsv2.31.1 (Quinlan and Hall 2010), using the CR1 BED file and respective assembly as input.

We aligned the extracted CR1 nucleotide sequences for each species to a reference sequence of the chicken (*Gallus gallus*, GenBank accession: U88211.1, accessed Sept 2021) complete consensus CR1 ORF2 using MAFFT (Katoh and Standley 2013). We used the ORF Finder tool in Geneious as an extra step to verify the boundaries of ORF2, and manually trimmed the CR1 sequences for each squamate species to the boundaries of the reference ORF2, followed by iterative realignment. After manual inspection, any individual sequences forcing large gaps (i.e., >500 bp) were removed from alignments.

We inferred species-specific CR1 phylogenies in IQ-Tree2 (Minh et al. 2020) with MFP and 1,000 bootstrap replicates (“-m MFP -B 1000”). The resulting 63 CR1 phylogenies were rooted using *G. gallus* as the outgroup and visualized in FigTree (Rambaut, 2010). Subclades were manually annotated based on cladistic groupings with the most recent common ancestor (MRCA) of each subclade with 100% parametric bootstrap. Sequences belonging to each subclade were then extracted from CR1 alignments using “seqtk” (Li 2013) ‘subseq’ to make concatenated multi-FASTA files for subclades from each species.

To verify that our CR1 sequences contained operable ORFs, we screened all sequences for the presence of a reverse transcriptase (RT) and ORF2 domain. We used usearch11.0.667 (Edgar 2010) with options “-fastx_findorfs-orfstyle 7-mincodons 16” to retrieve the ORF2 domain. We used hmmsearch (Finn et al. 2011) to generate a domain table for the RT domain against the Pfamv.28.0 database (Punta et al. 2012) as of May 2015 (includes 16,230 families) and collected all peptides annotated with RT. We then indexed the amino acid file via ‘esl-sfetch’, and generated a multi-FASTA file of identified RT hits using the index and RT domain table as input via ‘esl-sfetch’. Finally, we used faSomeRecords from UCSC Executables (Nassar et al. 2023) to generate new multi-FASTA files with sequences verified for presence of intact ORFs and RTs. Filtered, trimmed sequences were realigned using MAFFT and 50%-majority rule consensus sequences were inferred for each subfamily, resulting in a total of 118 lepidosaur CR1 subfamilies from 63 species. Sequences from 15 squamate species were filtered out for lack of intact ORF2 and RT domain (e.g., *Boaedon fuliginosus*, *Eryx tataricus*, see Supplementary Table 1).

### Phylogenomic Estimation of Squamate CR1 – Between-Species Analyses

To put CR1 evolution of squamate genomes in the context of amniote genome evolution as a whole, all available amniote CR1 consensus sequences were downloaded from RepBase v24 and aligned in Geneious Prime via MAFFT. The alignment was trimmed to target the ORF2 region and any sequences <1500 bp in length after trimming were filtered out. The amniote CR1 alignment was then filtered for intact ORF2 and RT domains using the scripts described above. A maximum-likelihood phylogeny was inferred using IQ-Tree2 with 1,000 nonparametric bootstrap replicates and the best-fit model determined by ModelFinder (Kalyaanamoorthy et al. 2017) (options “-m MFP-B 1000”) with all newly identified squamate CR1 subfamilies and amniote CR1 consensus sequences from RepBase.

### CR1 Genomic Distribution Analyses

To uncover the genomic distribution of squamate CR1 insertions, we downloaded protein sets for 20 squamate species available from RefSeq (O’Leary et al. 2016), Genome Warehouse, and Zenodo (last accessed December 2023). For the remaining 91 species without publicly available gene annotations, we generated annotation files using Liftoffv1.6.3 (Shumate and Salzberg 2021) with the annotation of each species’ closest relative (according to our species tree; see Supp Fig. 1). To ensure annotation quality, we generated proteome files using both LiftOff and original annotations via AGAT perl script agat_sp_extract_sequences.pl (option-p for amino acid sequences) (Dainat et al. 2023) and measured proteome completeness via BUSCOv 5.4.7 (OrthoDB v10 sauropsida database) in protein mode. We filtered all squamate proteomes by percent of complete BUSCOs > 80%, resulting in 30 accepted squamate genome annotations.

To understand the proximity of CR1 insertions to protein-coding regions, we combined our de novo repeat annotations with protein annotations for the 30 squamate species. We first parsed our protein annotations to isolate only exons and the largest isoform of each using custom bash scripts (see Data Availability Statement). We then used *bedtools closest* to measure the distance of each squamate CR1 element to the nearest exon (options-t all-D b), using the -D option to calculate distance with respect to upstream/downstream of exons. We also used *bedtools window* with *bedtools overlap* to identify CR1 elements overlapping exons (default settings).

For the purposes of investigating the CR1 genomic environment, we filtered CR1-exon distance results by CR1 length, with a minimum length of 500 bp and a maximum length of 5,000 bp to avoid falsely-identified repeats due to artifactual errors. To delineate between full-length and truncated CR1s, we separated results as follows: full-length CR1s > 2,500 bp (over 50% total length), truncated CR1s 500-2,500 bp (at least 10% total length). We used a phylogenetic generalized least-squares model to determine the impact of CR1-exon distances on CR1 length (generated using ‘nlme’ (Pinheiro et al.) and ‘ape’ (Paradis et al. 2004) R packages). We used Pagel’s lambda correlation structure with an initial lambda of 0.5 and a maximum-likelihood model fitting method. Using our species tree inferred for ancestral state reconstruction, we pruned to match our CR1-exon dataset, dropping one species absent from our tree, resulting in 29 species. To account for variable intergenic distances across squamate species, we also calculated intergenic distances for 29 squamate genomes using custom python scripts and generated a PGLS model with both CR1 length and intergenic distance as response variables with the same settings.

Lastly, we calculated GC content for 1kb flanking regions downstream and upstream of all CR1 inserts, filtered by length. We used seqkit subseq with flags ‘-u’ and ‘-d’ along with ‘-f’ and our de novo repeat annotations to generate flanking regions, followed by seqkit fx2tab with option ‘--gc’ to calculate GC% for each flanking region.

## Data Availability Statement

Whole genome files, alignments, tree files, and repeat annotations have been made publicly available at NCBI (PRJNA1073866; PRJNA1073867) and Zenodo at https://doi.org/10.5281/zenodo.10521011.

## Acknowledgements

We would like to acknowledge the Office of Undergraduate Research and Creative Activity and the Monsoon High Performance Computing Cluster at Northern Arizona University. This work was supported by the National Science Foundation under grant DEB 2323124 to MT, a State of Arizona Technology Research Initiative Fund (TRIF) Faculty Support Grant awarded to MT, and in part through the National Institutes of Health U54 CA217376. Unpublished genome assemblies for *Uta stansburiana* and *Intellagama lesueurii* are used with permission from the DNA Zoo Consortium (dnazoo.org). The authors thank Emma Reich, Panth Patel, and Vahid Fard for feedback on this manuscript.

