## Supplementary Materials for "Differential Conservation and Loss of CR1 Retrotransposons in Squamates Reveals Lineage-Specific Genome Dynamics across Reptiles"

**Supplementary Figures**

Simone M. Gable[1], Nicholas Bushroe[1], Jasmine Mendez[1], Adam Wilson[1], Brendan Pinto[2], Tony Gamble[3], Marc Tollis[1*]

***Supplementary Figure 1:*** *Squamate coalescent-based species tree inferred from gene trees with branch support shown in posterior probabilities on all nodes.*

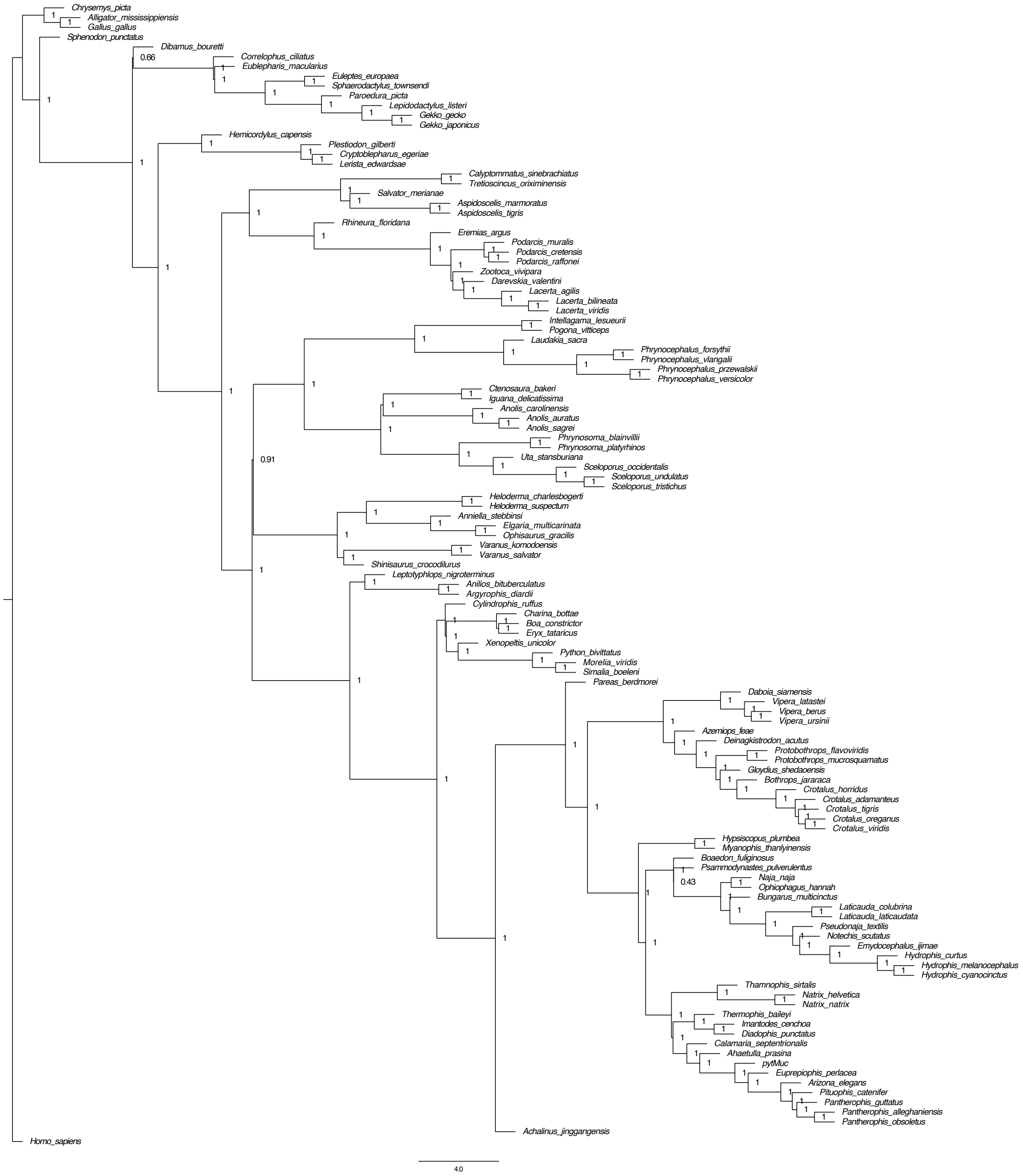

***Supplementary Figure 2:*** *Turtle coalescent-based species tree inferred from gene trees with full branch support shown in posterior probabilities on all nodes.*

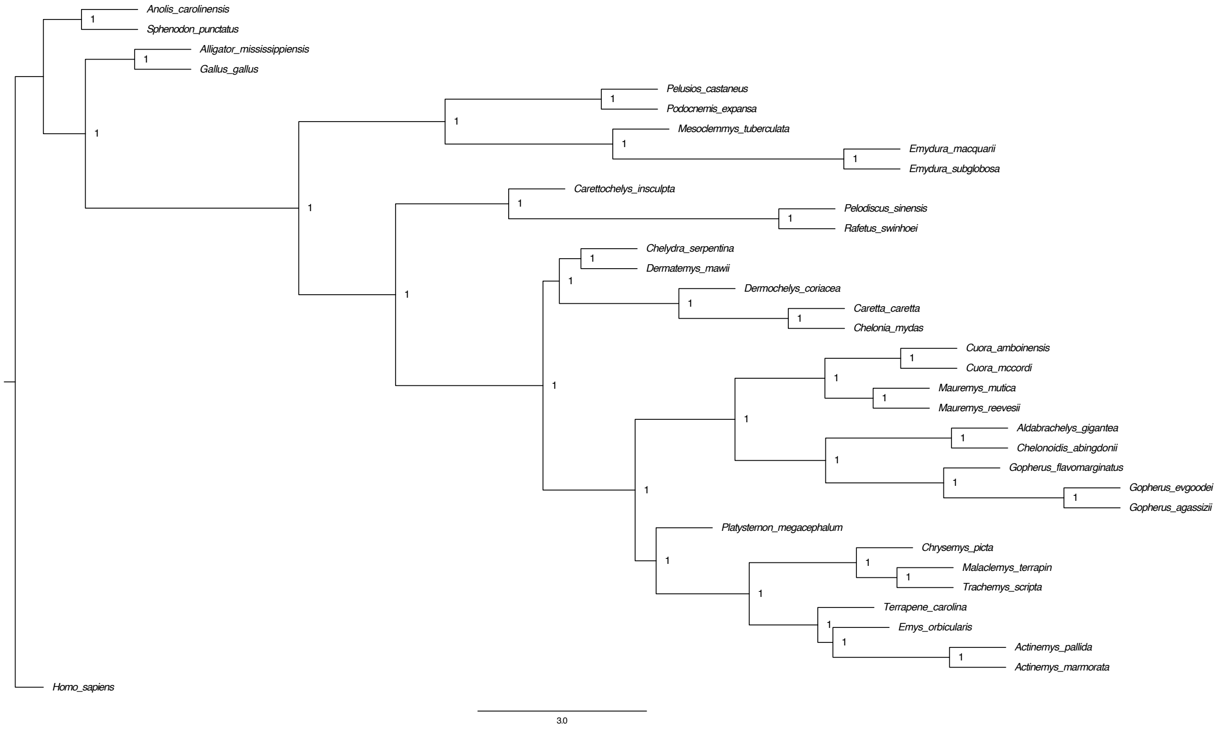

***Supplementary Figure 3:*** *Archosaur coalescent-based species tree inferred from gene trees with branch support shown in posterior probabilities on all nodes.*

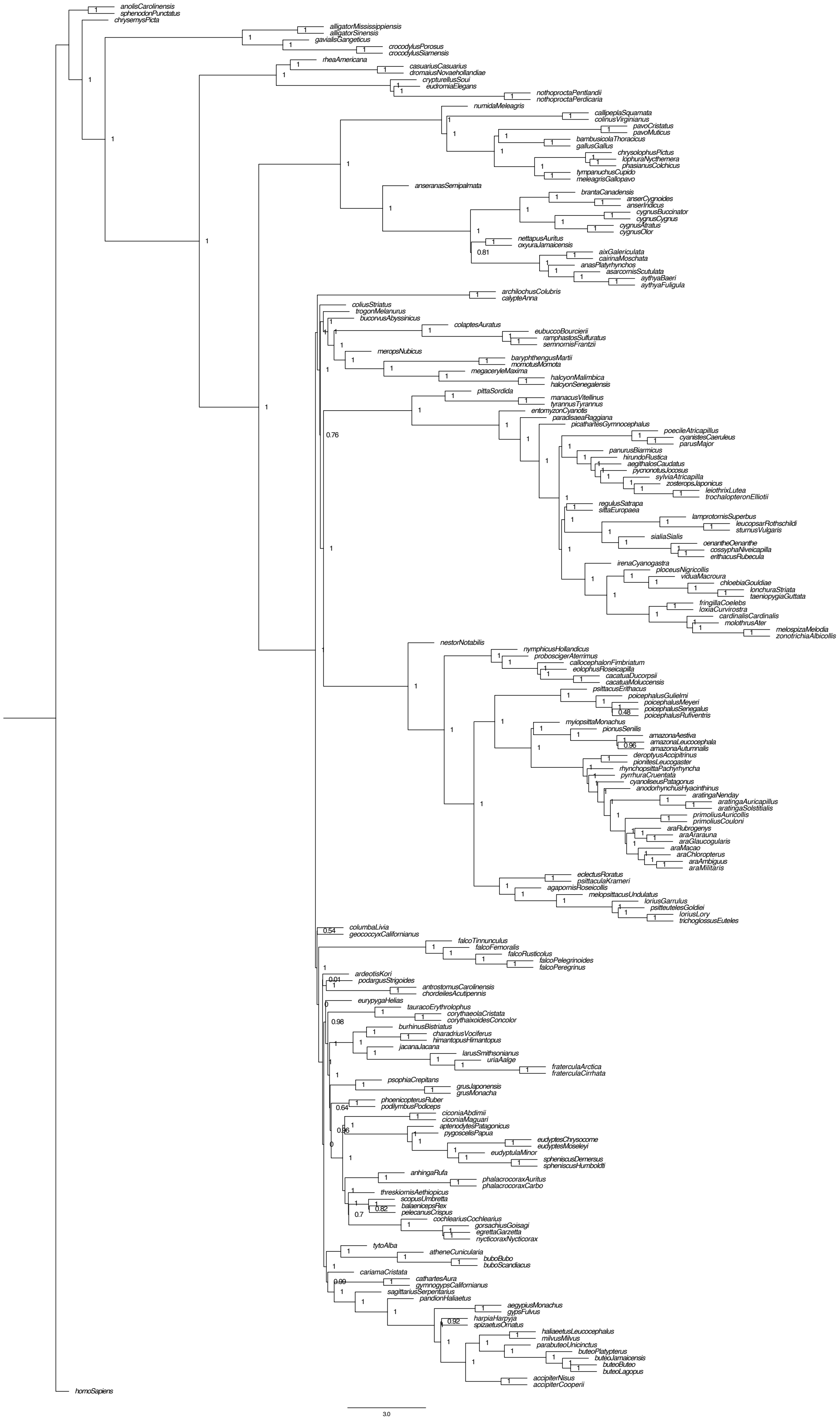

***Supplementary Figure 4:*** *CR1 repeat landscapes of species with highest amount of significant Wilcoxon Pairwise comparisons (p<0.05), as well as A. carolinensis for reference.*

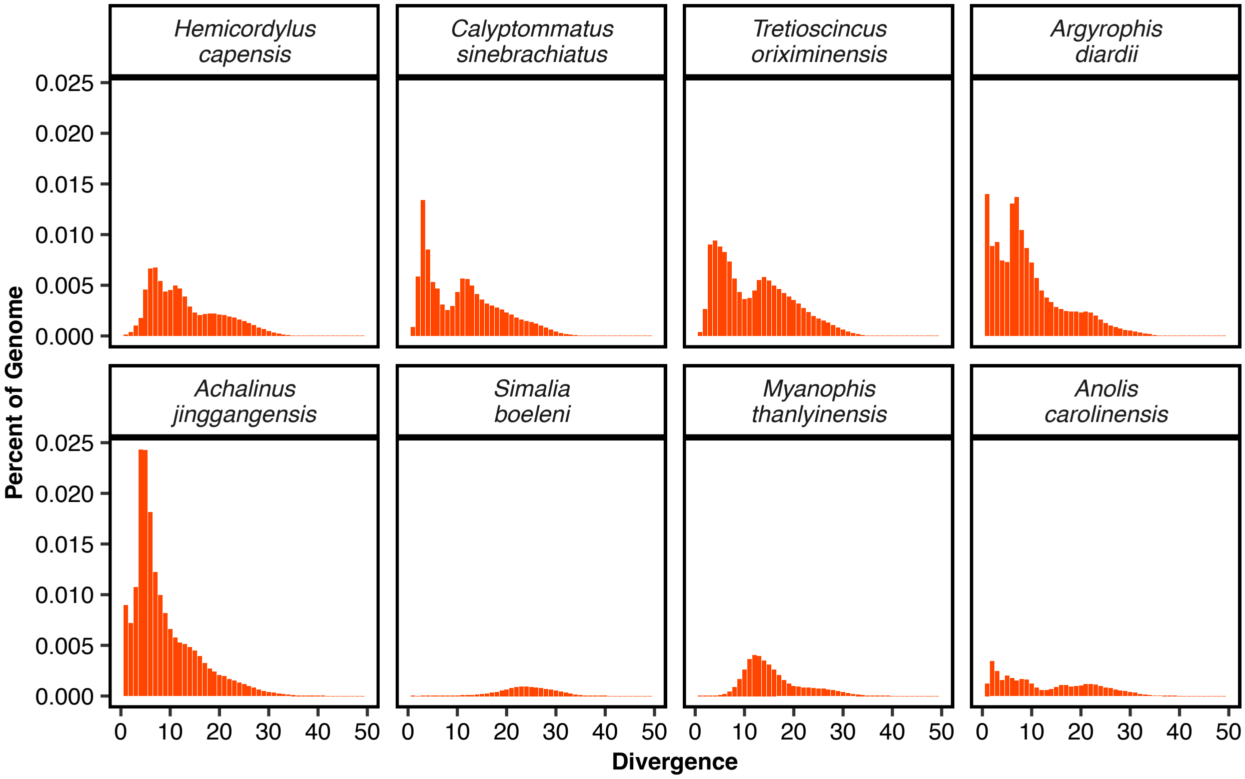

***Supplementary Figure 5****: Distributions of CR1 copy number for squamate (green), turtle (blue), and archosaur (purple) species. Bin width =  20,000.*

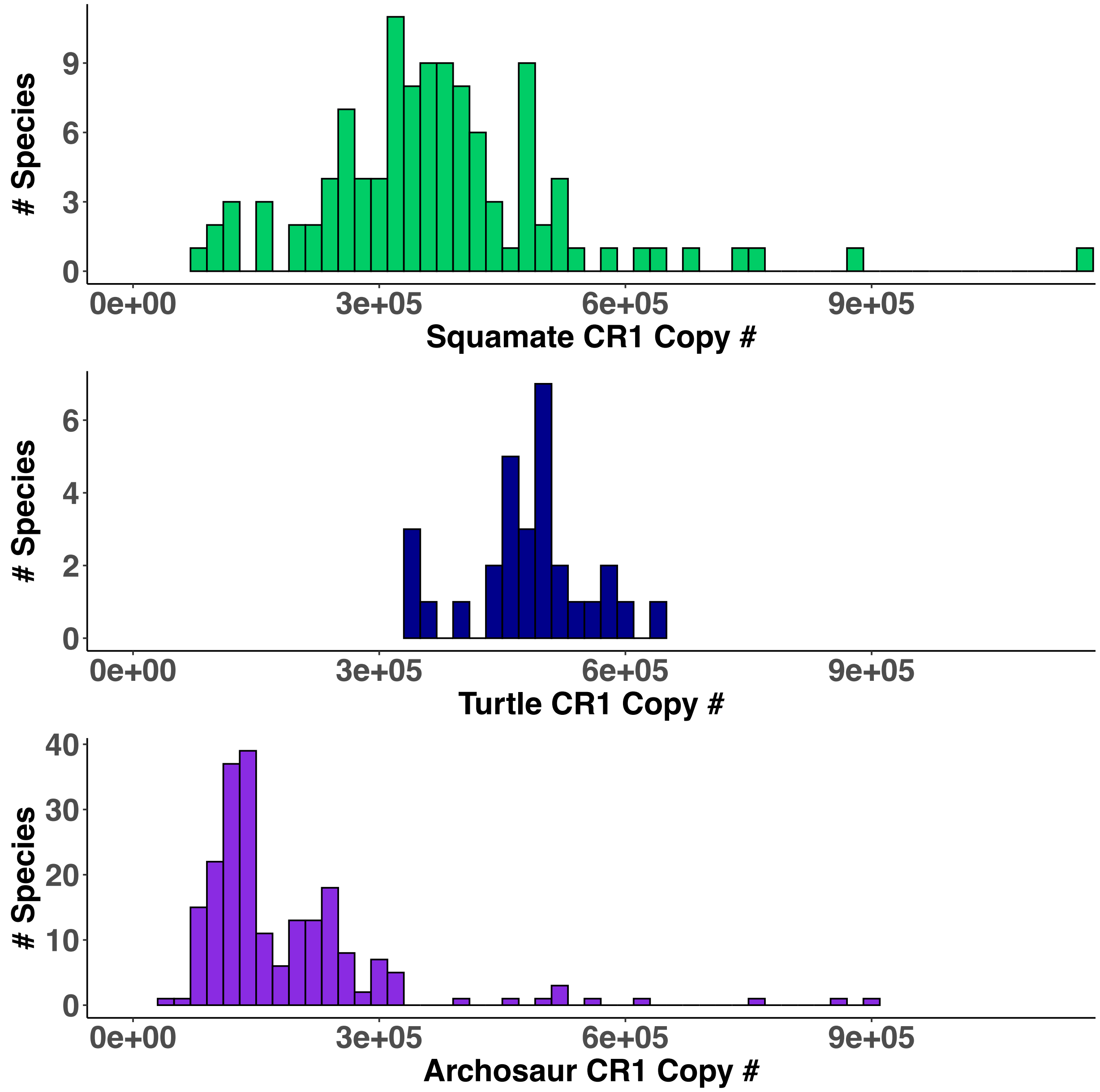

***Supplementary Figure 6:*** *Linear regression of CR1 copy number vs. assembly size (b.p.) for 111 squamate genomes.*

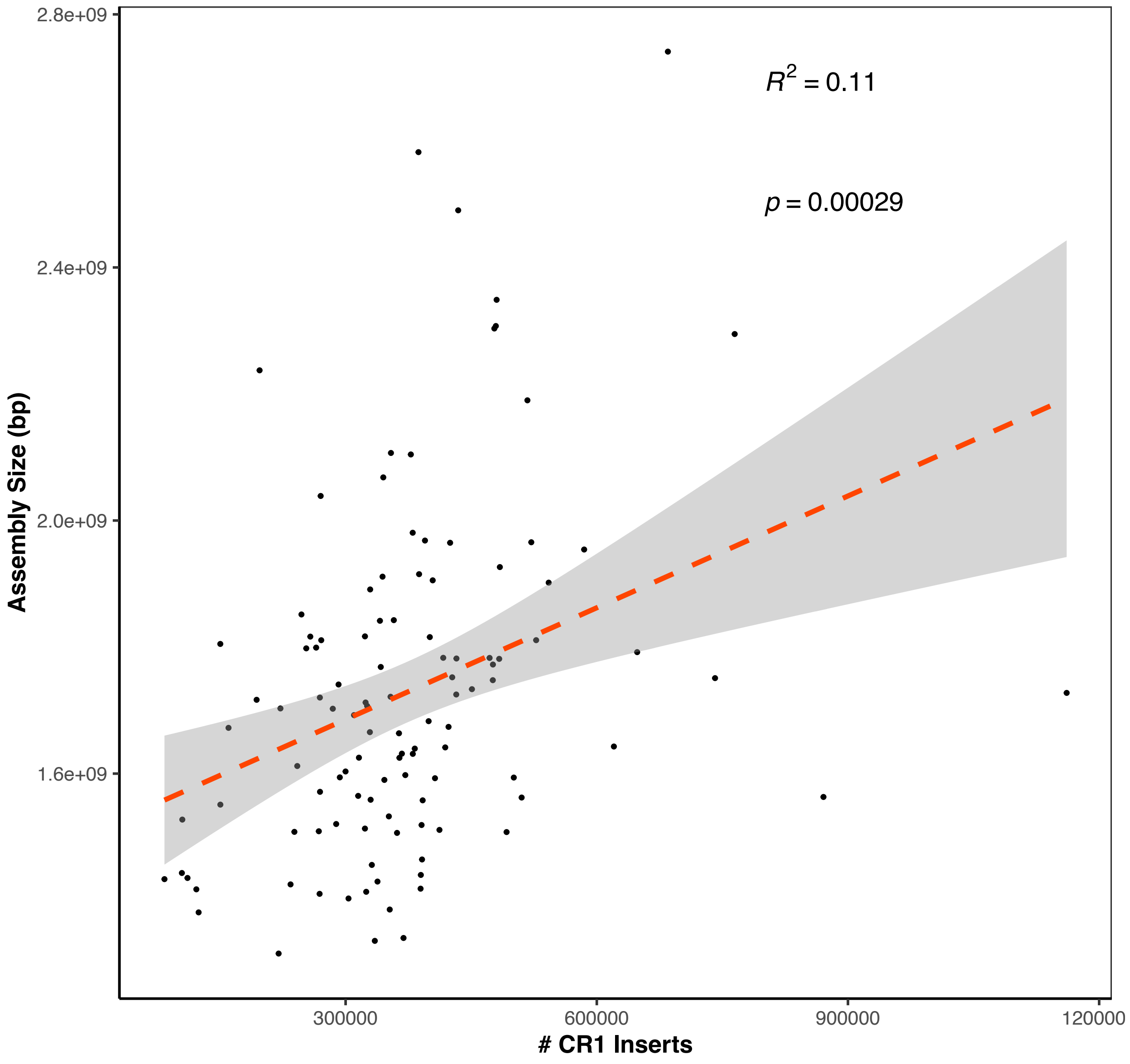

***Supplementary Figure 7:*** *QQ-Plots for CR1 copy number rates for turtles, squamates, archosaurs, and birds. Blue lines represent slope of line connecting points at quartiles of theoretical and sample distributions. Turtle rates are normally distributed, while squamate, archosaur, and avian-only (bird) rates are non-normal.*

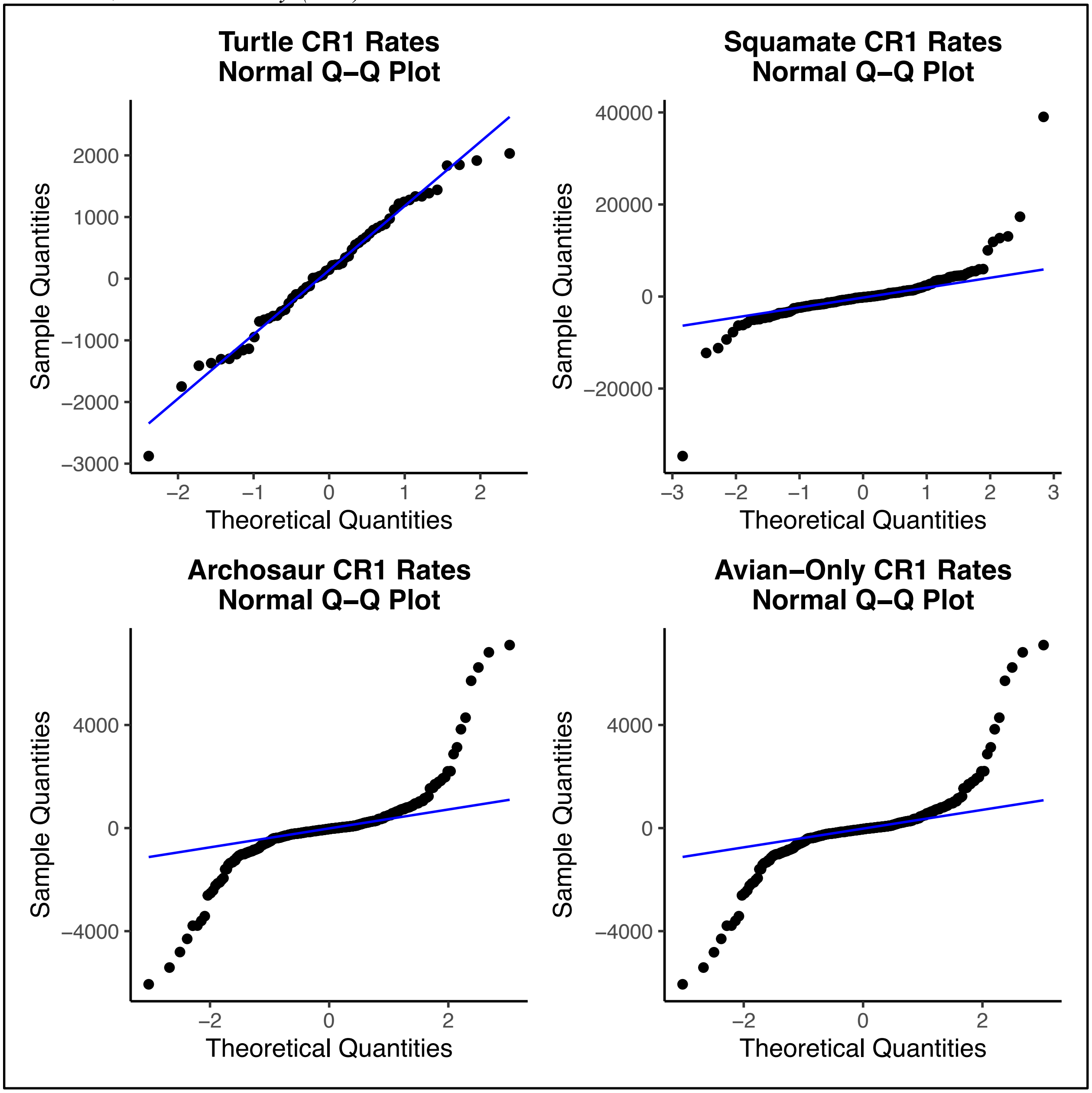

***Supplementary Figure 8:*** *Ultrametric squamate tree inferred using coalescent-based tree with dates estimated in TreePL. Geological timeline from TimeTree.org (see Chapter 3 Methods "Divergence Time Estimation").*

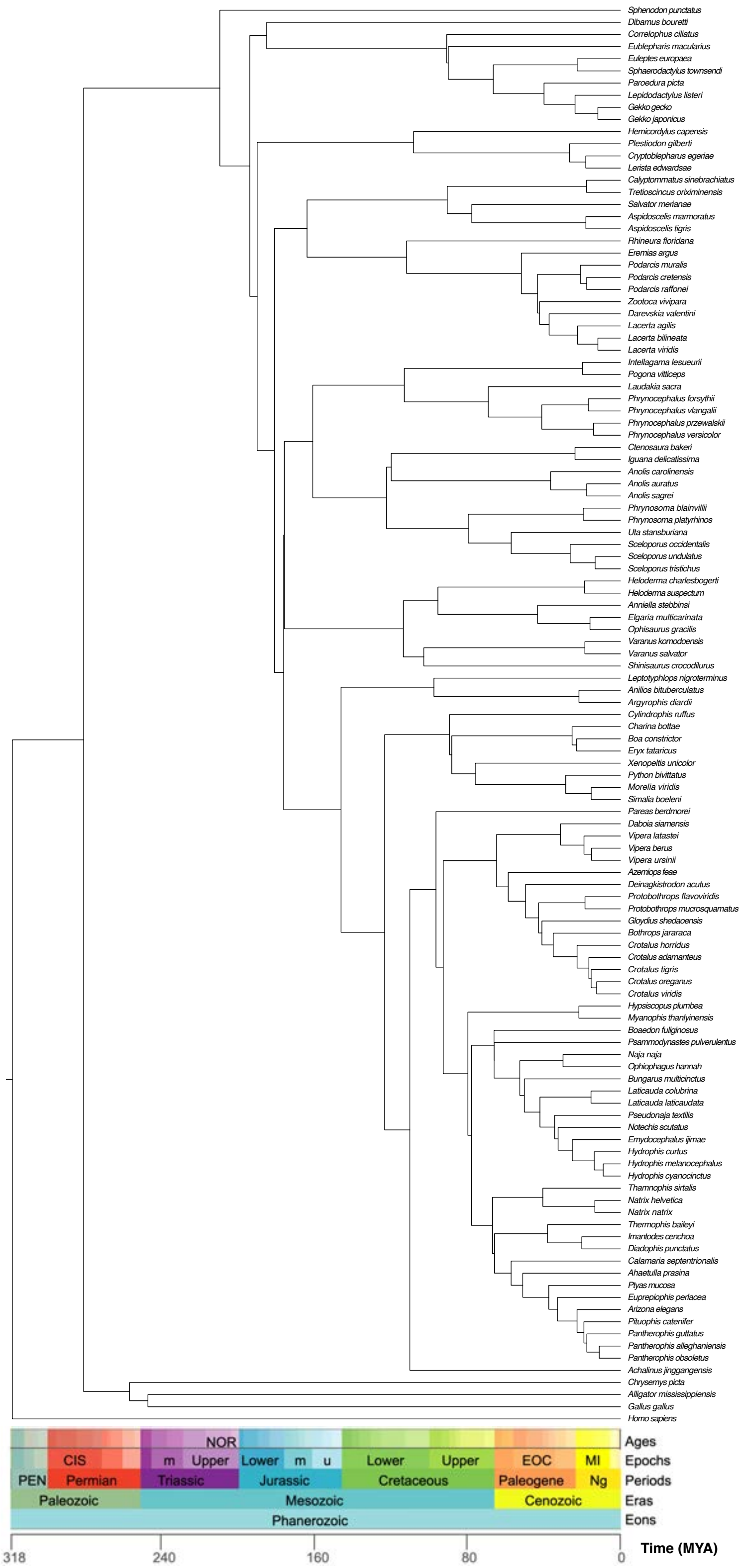

***Supplementary Figure 9a:*** *Turtle species tree 3,342 orthologs nucleotide sequences of high-confidence single-copy orthologs with ancestral state estimation mapped onto branches. Branch lengths are in terms of coalescent units. Rooted using H. sapiens outgroup (not shown). CR1 insert number was modeled as a continuous trait for all genomes with annotated repeats.*

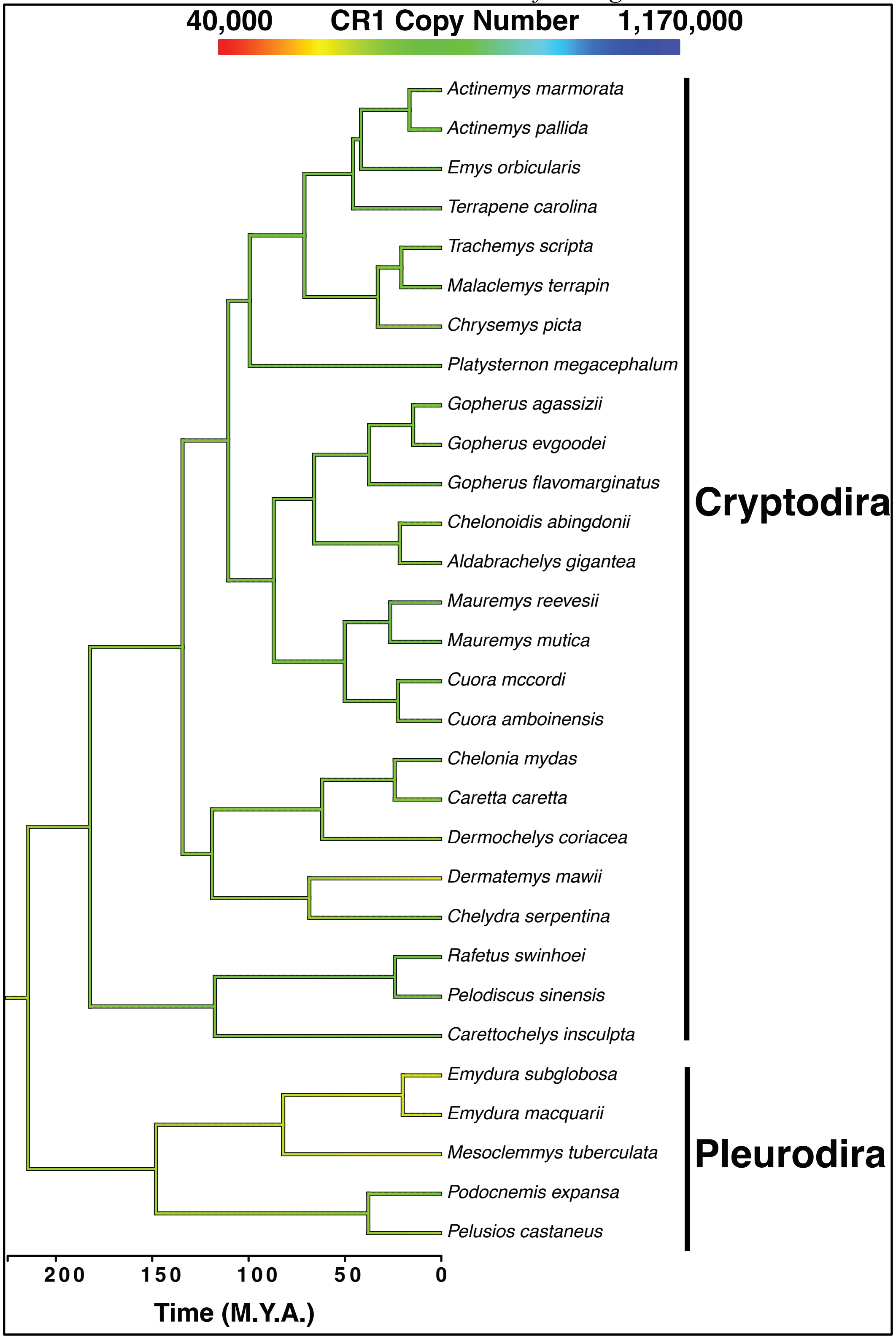

***Supplementary Figure 9b:*** *Archosaur species tree constructed with 2,981 nucleotide sequences of high-confidence single-copy orthologs with ancestral state estimation mapped onto branches. Branch lengths are in terms of coalescent units. Rooted using H. sapiens outgroup (not shown). CR1 insert number was modeled as a continuous trait for all genomes with annotated repeats.*

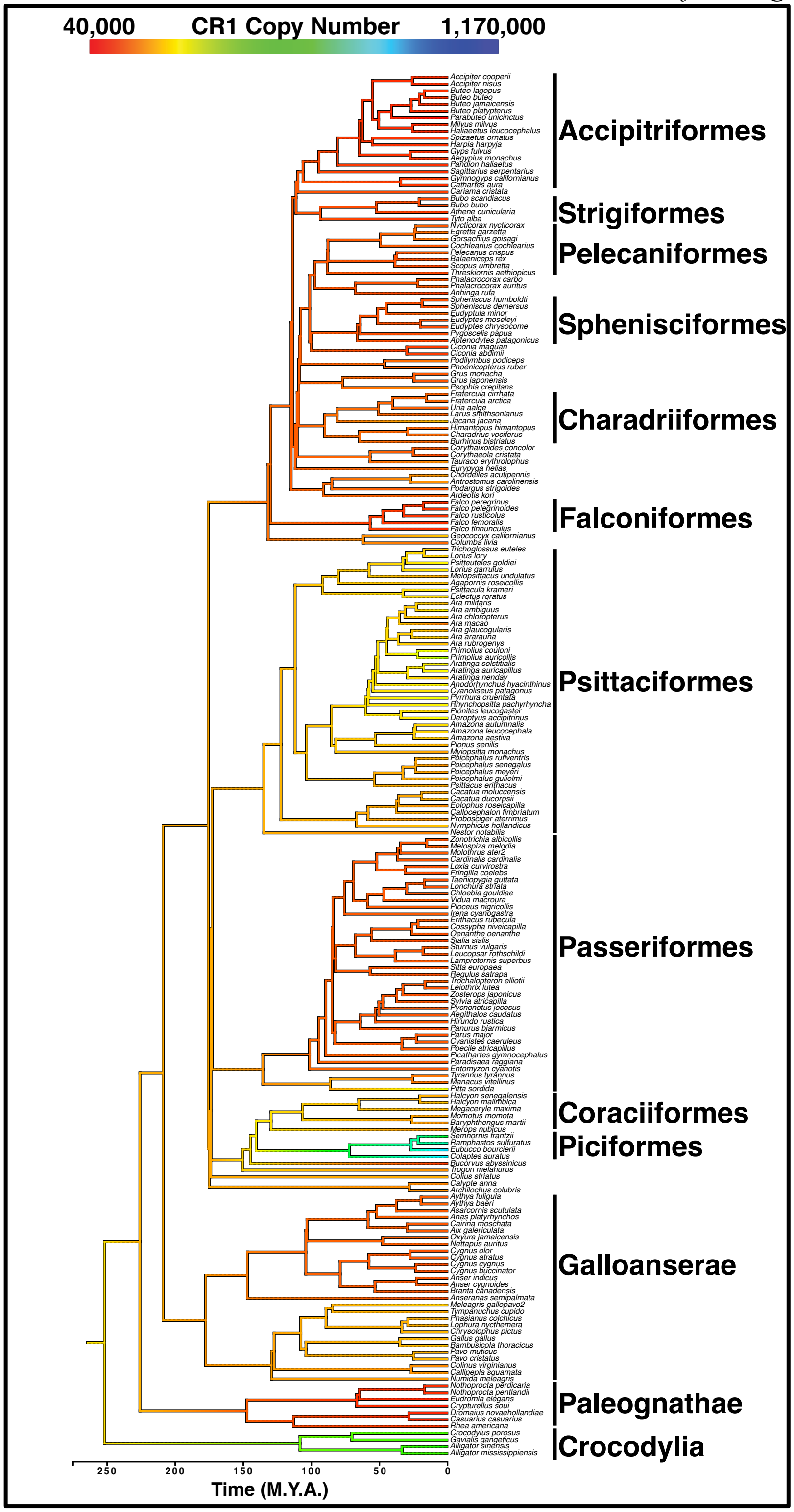

**Supplementary Figure 10:** Scatterplots for 30 squamate genomes comparing distance of each CR1 to nearest exon (kbp) vs. CR1 length (bp).

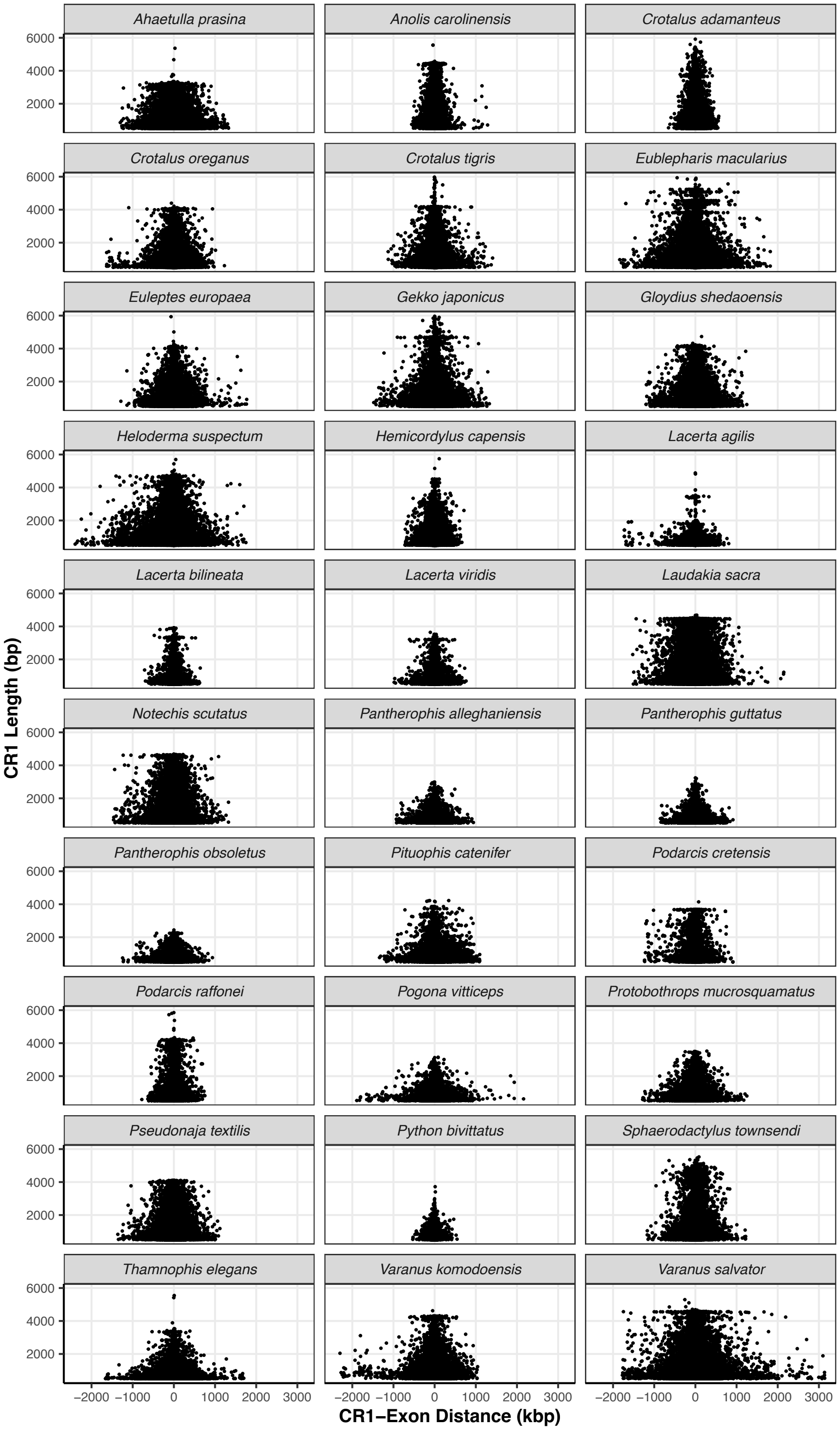
